# Glycation renders ɑ-synuclein oligomeric strain and modulates microglia activation

**DOI:** 10.1101/2022.01.15.476311

**Authors:** Manisha Kumari, Bhoj Kumar, Krishna Singh Bisht, Tushar Kanti Maiti

## Abstract

*α*-Synuclein is known to involve in the pathogenesis of Parkinson’s diseases (PD) and related disorders. However, it is unclear how its aggregation causes neuronal degeneration and neuroinflammation. Due to intrinsic disorder nature, *α*-synuclein produces a large number of structural ensembles and diverse aggregation intermediates. The post-translational modifications add a new layer of complexity to the aggregation mechanism. Recently, it has been demonstrated that glycation of *α*-synuclein restricts into oligomeric intermediates and causes neuronal toxicity. However, the understanding of aggregation mechanism, dopaminergic neuronal death, and neuroinflammation by the glycated *α*-synuclein is yet to be elucidated. The present study aims to address how glycated synuclein differs in oligomerization and neuroinflammation. The glycation of *α*-synuclein perturbs the aggregation kinetics and prevents the fibrilization through the alteration of surface charges of N-terminal domain residues which prevents membrane binding and seed amplification mechanism. Mass spectrometry-based proteomics analysis of BV2 cells treated with glycated oligomers provides evidence of alteration of endocytic mechanism, mitochondrial dysfunction, and inflammatory cascade. Here, we show that *α*-synuclein oligomers strongly bind to TLR2 and activate the TLR2 mediated signaling. However, glycated *α*-synuclein oligomers impair the TLR2 binding and compromise TLR2 signaling. Interestingly, we also find that the glycated *α*-synuclein oligomers favor NLRP3 inflammasome mediated neuroinflammation compared to non- glycated *α*-synuclein oligomers. In conclusion, our findings suggest that microglia response towards *α*-synuclein is conformation-specific and glycated oligomers can contribute to neurodegeneration differently.

## Introduction

Parkinson’s disease is the one of common neurodegenerative disorder worldwide which affects almost 2% of the world population over the age of 60 [1]. The pathological hallmark of PD is associated with the progressive loss of dopaminergic neurons in the substantia nigra along with chronic inflammation, mitochondrial dysfunction and accumulation of *α*-synuclein rich protein aggregates in the form of Lewy body [2]. α-Synuclein is an intrinsically disorder protein (IDP) with a molecular weight of ∼14.5 kDa that plays a crucial role in synaptic vesicle formation and its trafficking [3], neuronal differentiation [4], suppression of apoptosis [5], regulation of dopamine biosynthesis [6]. α-Synuclein carries three main domains where N-terminal (1-60 residues) is comprised of positively charged amino acids and facilitates membrane binding, the middle NAC region (61-95 residues) bears majorly hydrophobic residues that are responsible for amyloid fibre formation and the C-terminal (96-140 residues) is majorly consists of acidic amino acids which binds with calcium and promotes it’s entry to the neuronal cells. C-terminal domain also plays a pivotal role in protecting NAC domain aggregation [7]. Interestingly, 140 amino acid residues of *α*- synuclein provides hydrophobic and electrostatic intermolecular interactions which control the solubility of the monomeric form [8]. These intermolecular interactions are vastly influenced by the cellular milieu and imperceptible change can lead to slant of the α-synuclein conformation [9]. Thus, destabilization of the monomeric α-synuclein conformation favours aggregation and generate toxic species [11].

Post-translational modifications such as phosphorylation [12], O-GlcNAcylation [13], nitration [14], acetylation [15], ubiquitination [16], SUMOylation [17] and glycation [18] play an important role in its structure, function and aggregation. Recent studies have demonstrated that there is close association of type 2 diabetes mellitus and Parkinson’s disease. Individuals with diabetes mellitus have 35% risk of developing cognitive and motor impairment and finally Parkinson’s disease [19]. In fact, hyperglycaemic state in an individual produces more methylglyoxal (MGO) that is produced due to altered carbohydrate metabolism or impairment of glyoxalase system [20]. MGO is a di-carbonyl compound that spontaneously modifies Lysine, Arginine and Cysteine residues of protein and generates advance glycation end products (AGEs) [21]. Proteins with AGEs have been shown at the periphery of the Lewy body and their levels are increased in the PD patient brain [22]. It has been documented that *α*- synuclein undergoes MGO modification majorly in its N-terminal residues and inhibits membrane binding, promotes the accumulation of toxic oligomers and inhibits neuronal synaptic transmission [23]. MGO modification α-synuclein also reduces its binding to Cu^2+^ metal hence abolishes the free radical scavenging property of α-synuclein [24]. Overexpression of chaperones such as DJ-1 and Hsp27 mitigates the glycation induced aggregation and cytotoxicity [25, 26]. It is interesting to note that glycation of α-synuclein restricts itself into oligomeric conformation and inhibits amyloid fibril formation [23]. The molecular mechanism of oligomeric restriction and neurotoxicity has yet to be unravelled and detail mechanistic study is necessary.

The microglial activation leads to chronic neuroinflammation and it is also one of the hallmarks of PD [27]. An activated microglial signature has been observed in the post-mortem PD brain [28]. Microglia tirelessly surveils the micro-environment and able to recognize a minute change [27, 29]. During neuronal transmission of α-synuclein, microglia senses, engulfs and clears the extracellular α-synuclein to maintain the homeostasis in the brain (30, 31). Several microglial receptors such as TLR2, TLR4, CD11b, FcγRIIB have been reported to interact with α- synuclein and its strains further induce pro-inflammatory microglia response [32, 33, 34, 35]. The uptake of α-synuclein oligomeric species is dependent on TLR2 receptor of microglia [36]. TLR2 mediated microglia activation is conformation specific, monomeric and fibrillar species were fail to act as TLR2 agonist [36]. Recently, NLRP3 inflammasome activation has been reported in post-mortem PD brains [37, 38]. Localization of both NLRP3 and ASC was also confirmed in activated microglia in PD patients [39]. Interestingly, α-synuclein fibrils have also been shown to activate the NLRP3 inflammasome [40]. α-Synuclein produces large number of structural ensembles and produces diverse oligomeric intermediates, it is conceivable to believe that these intermediates evoke differential neuroinflammation. This brings up a question that how does these diversified ɑ-synuclein species and strains modulate microglia inflammatory processes. As MGO modification to α-synuclein locks the oligomeric structure and contributes higher neuronal toxicity and it is a pertinent question for us to check if MGO modified oligomeric species show differential neuroinflammation compared to unmodified oligomeric species.

In this study, we used various biophysical and biochemical methods to demonstrate the conformational differences between glycated synuclein oligomers (MO) and wild type synuclein oligomers (SO), along with their conformation driven microglia activation pathways. We used the abbreviation MO and SO throughout the manuscript. Our results show that glycation of α-synuclein generate oligomers of different morphology and exhibits reduced membrane binding. The surface charge alteration in the N-terminal domain inhibits seed amplification for further fibrilization of α-synuclein. Our findings also reveal that MO choose clathrin dependent endocytic pathway and activate microglia preferably via inflammasome pathway. Holistically, this study addressed the link between the diversity of synuclein oligomeric strain and the modulation of neuroinflammation.

## Results

### Glycation generates oligomers with different morphology and secondary structures

Though methylglyoxal modification of α-synuclein has been reported earlier, its impact on structure and aggregation mechanism has not been clearly elucidated. Earlier literatures describe that MGO modification restricts α-synuclein into oligomeric conformation during aggregation (23). However, molecular basis of this restricted oligomeric structure formation has not been established. Here, we generated glycated α-synuclein by incubating 200 μM α- synuclein with 5 mM MGO at 37 ^0^C for 24h (Figure 1A). We characterized the MGO modified species using mass spectrometry (MS), atomic force microscopy (AFM), Transmission electron microscopy (TEM), CD spectroscopy and dynamic light scattering (DLS). The intact mass analysis of glycated α-synuclein by MALDI-MS confirmed an increase of 389 Da molecular mass upon addition of MGO (Figure 1B). Molecular weight of low molecular weight species (LMW) of α-synuclein was found to be 14.431 kDa, whereas upon glycation its molecular weight became 14.82 kDa. Further using trypsin digestion followed by mass spectrometry, we mapped the MGO modification sites. We found multiple modified lysine sites with high and low identification score, out of which we selected lysine with high identification score. Lysine residue of 21, 23, 45, 58, 60, 96, 97 positions (Supplementary Figure 1) were modified and these sites were consistent with the earlier reported results. These lysine adducts are largely reside in N-terminal domain. Next, we examined the aggregation kinetics of LMW α-synuclein under MGO treatment condition via Thioflavin (ThT) dye. In presence of MGO, α-synuclein showed weak binding to ThT during the aggregation process. However, untreated LMW α-synuclein showed sigmoidal ThT binding kinetics which comprised of lag phase, exponential phase and stationary phase (Figure 1C). Atomic force microscopy image was taken at 52 h time point of aggregation kinetics and revealed that MGO modified species stayed in the oligomeric conformation. However, unmodified LMW α- synuclein formed fibrils with defined periodicity (Figure 1D).

**Figure 1:**
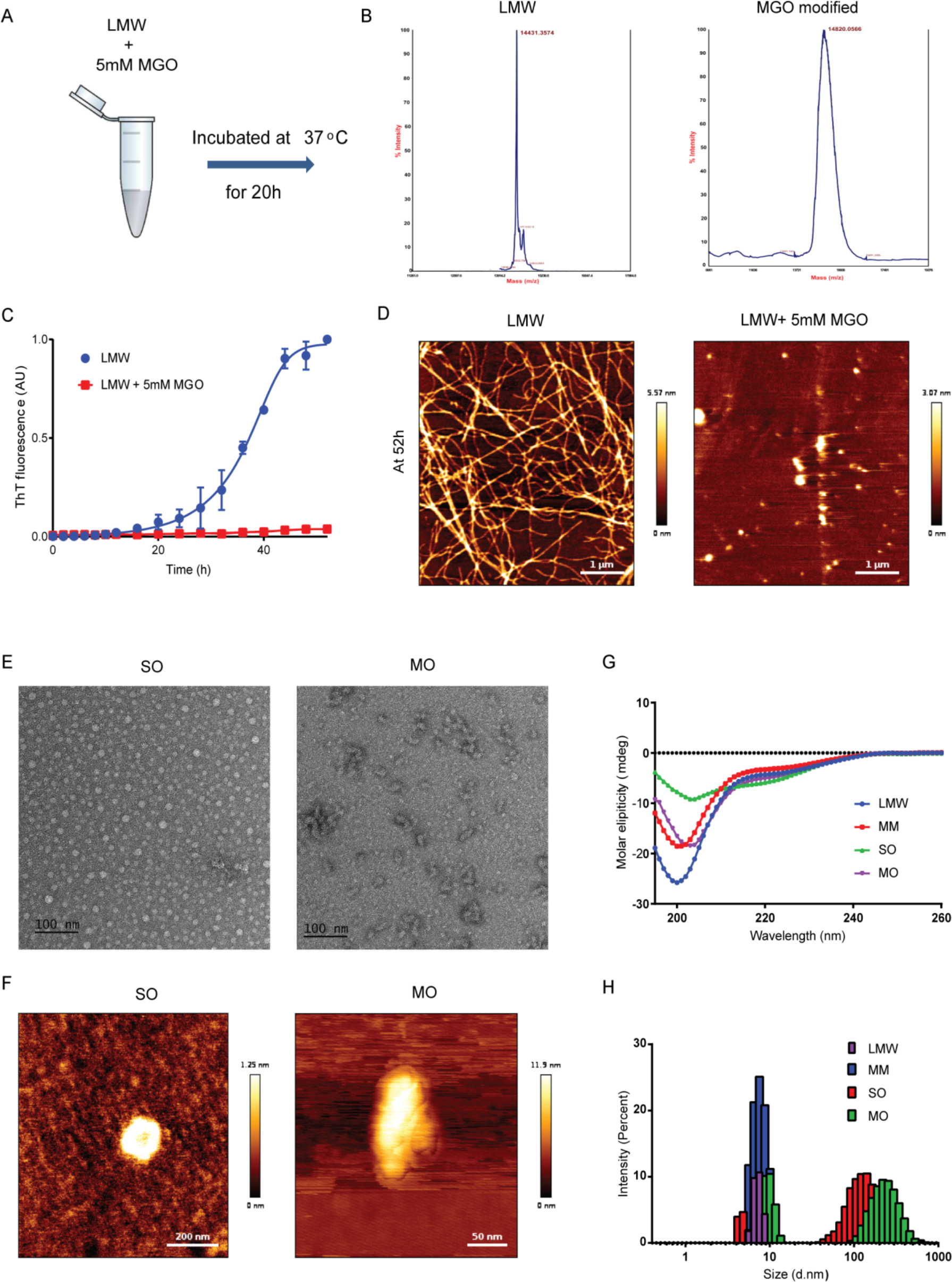
Characterization of MGO modified synuclein. (A) Schematic representation of preparation of glycated α-synuclein. LMW synuclein species were incubated with 5 mM MGO at 37 ͦ C for 20 h. (B) Intact mass analysis using MALDI mass spectrometry demonstrating increase of mass of modified synuclein. (C) ThT fluorescence measurement representing aggregation of LMW and MM species, n=3. (D) AFM images showing morphology of aggregates at 52h, LMW acquired fibrillar end state, whereas MGO modification restricts synuclein in oligomeric state. (E, F) High resolution TEM and AFM showed distinct morphology synuclein oligomers (SO) and Modified oligomers (MO), respectively. (G) CD spectroscopy revealed secondary structure of SO and MO, n=3. (H) Size distribution analysis using DLS showed MO having pool of different size of oligomers than SO, n=4.

Due to natively unfolded structure, α-synuclein can adopt different oligomeric conformations assorting from spherical, annular, donut shape/pore like, which drive the neuronal toxicity differentially. So, we aim to understand the morphological differences between α-synuclein oligomers (SO) and MGO modified oligomers (MO). We performed the high resolution TEM imaging along with high resolution atomic force microscopy (AFM) to dissect the morphology of both unmodified and modified species (Figure 1E and 1F). α-Synuclein oligomers showed donut shape/ring like in morphology with the height of 10-14 nm, whereas, MGO modified oligomers comprised of both elongated and amorphous morphology combining 2-4 isolated units with height of 8-14 nm. LMW and MGO modified LMW (MM) showed random coil secondary structures as monitored in CD spectroscopy (Figure 1G). MGO modification did not affect the secondary structure of LMW α-synuclein. However, oligomeric species generated from MGO modified α-synuclein (MO) showed significant lower amount of beta sheet than SO. Further, Dynamic light scattering (DLS) indicated that the diameter of MO was more than SO (Figure 1H). Taken together, our observations indicate that MO has different morphology than SO and carries less beta sheet secondary structure upon oligomerization.

### Glycation of synuclein decrease the secondary nucleation and lipid binding by altering the exposed surface charges and hydrophobic patches

We speculate that MGO modified α-synuclein must follow the different kinetic pathways in the aggregation process and rearranges tertiary structure that restricts the conversion into beta sheet. Secondary nucleation and membrane binding of α-synuclein dependent on the solvent exposed charged residues and hydrophobic surface (43, 44). So, to understand the overall topology of glycated and unmodified species of α-synuclein, we first calculated the surface charges of all species. We found that LMW α-synuclein showed zeta potential -14 mV which represents the surface charge contribution of disordered species. MGO generates a negative charge on lysine residue upon modification. Thus, we saw the change of Zeta potential to -25 mV (Figure 2A). In case synuclein oligomer (SO) the zeta potential further shifted significantly than LMW due to compact structure formation by positively charged reside and flanking C-terminal negatively charged residues (45). We observed highest negative zeta potential in MO species (-29 mV), possibly due to modification of solvent exposed N-terminal lysine residues (Figure 2A). The Zeta potential data suggested that the oligomeric species generated in glycated synuclein and non-glycated synuclein share different molecular arrangement. The zeta potential data is also substantiated by ANS binding experiments of all the α-synuclein species. ANS binds to the hydrophobic surface and generates the fluorescence emission peak near 515- 525 nm. LMW species showed emission maxima around 510 nm, however, the intensity was less. In case of SO species, ANS strongly binds and showed emission maxima 500 nm. On the other hand MO showed strong fluorescence intensity with highest blue shift (emission maxima 450 nm). Thus ANS binding demonstrated that hydrophobic surface was exposed more in MO and MM species than the SO and LMW, respectively (Figure 2B). This data reflect that glycation modulating overall topology of these species.

**Figure 2:**
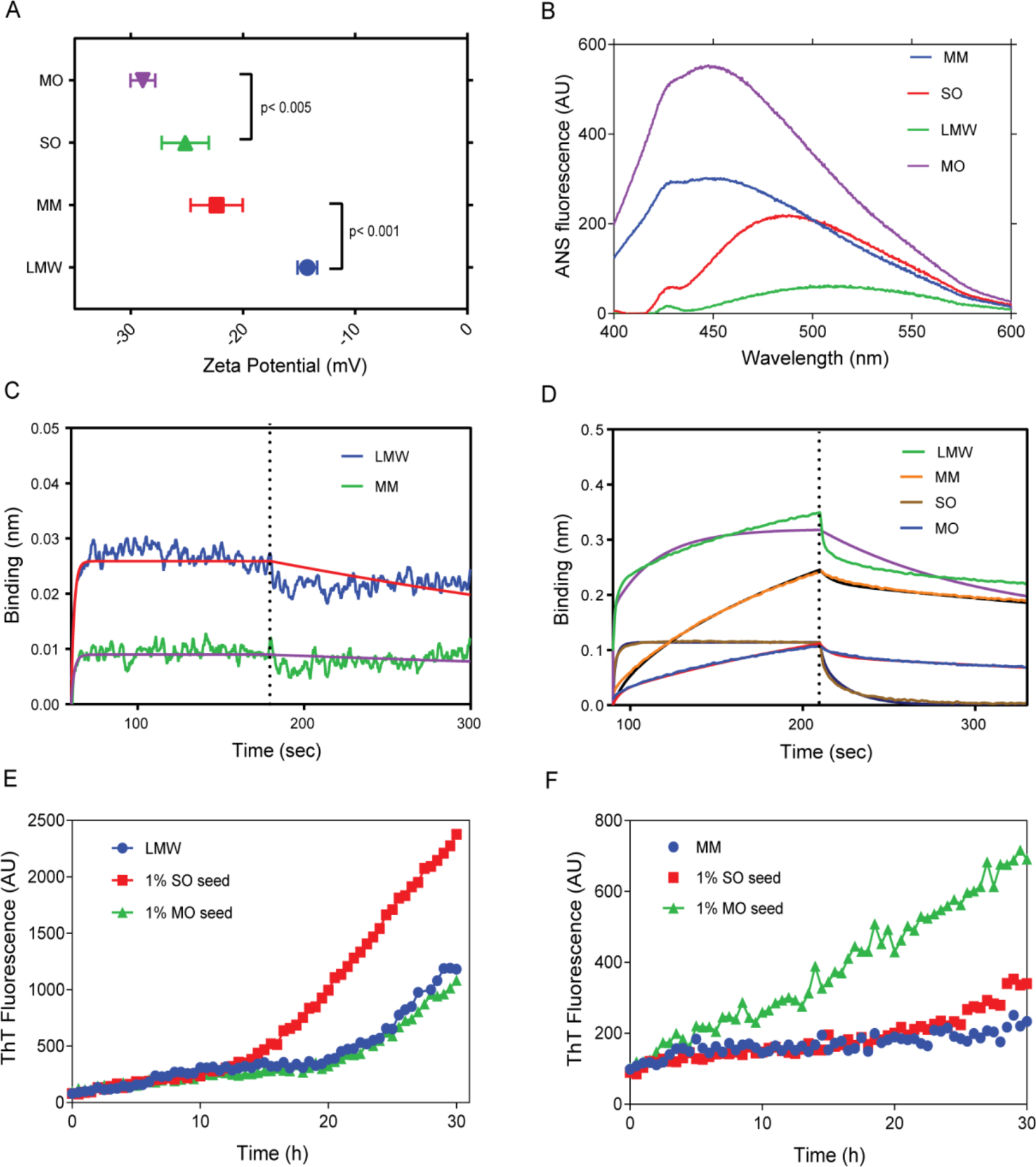
Glycation modulate surface charges and hydrophobicity of α-synculein. (A) Zeta potential analysis revealed the overall surface charges of synuclein species. MO and MM showed more negative potential than SO and LMW, respectively, n=3. Statistical significance was calculated using one way ANOVA followed by Tukey’s multiple comparison test in GraphPad Prism software. (B) ANS fluorescence upon excitation at 340 nm revealed that MO and MM expose greater hydrophobic surface to solvent than SO and LMW, n=3. (C) Biolayer interferometry (BLI) graph demonstrating binding of lipid with LMW and MM species of α- synuclein, were measured by immobilizing LMW and MM over AR2G sensors tips and dipped in SUVs contacting well. (D) BLI graph showing the interaction of α-synuclein species with LMW synuclein by loading LMW species over AR2G sensor tips and dipped in wells containing 10 μM of α-synuclein species. Among all species, SO interacted significantly more to AR2G sensor loaded LMW. (E) ThT fluorescence demonstrating aggregation kinetics of 50 μM of LMW incubated with 1% seed of SO and MO, n=3. (F) ThT fluorescence demonstrating aggregation kinetics of 50 μM of MM incubated with 1% seed of SO and MO, n=3.

MGO modified several lysine residues majorly in the N-terminal region. It has been documented that N-terminal surface lysine residues play a crucial role in stabilizing the α- synuclein binding with the lipid/neuronal membrane. To understand the role of glycation in membrane binding, we performed Biolayer interferometry (BLI), where we measured the binding of MM and LMW with the DOPC:DOPE:DOPS lipid membrane. LMW species recruited rapidly and immediately reached to the saturation binding with KD= 9.66E-07±5.9 M. On the other hand MM species showed slow binding and KD was found to be 2.42E-03±0.31 M (Supplementary Table 1). As MGO modified surface lysine residues at the N-terminal and generated an overall negative charge to the surface, we observed decreased membrane binding in case of MM compared to the LMW (Figure 2C).

Next, we explored to find the link between the morphology of oligomers and seed amplification mechanism. Seed amplification mechanism has been well illustrated in the “on-pathway” of amyloid protein aggregation. The amplification of seed towards protofibril formation proceeds through recruitment of monomeric α-synuclein on the surface of oligomeric seed and this is one of the key events in the fibrilization process. To shed a light on seeded aggregation, we performed the BLI experiment and analysed the binding of α-synuclein LMW species with LMW, MM, SO and MO species. We loaded the AR2G sensors with LMW species and dipped in the wells containing different species of α-synuclein. We found both MM and MO species showed decrease in binding towards sensor loaded with LMW in comparison to LMW and SO species (Figure 2D, Supplementary Table 2). Further, to understand secondary nucleation capacity of oligomers, we studied the aggregation kinetics of LMW and MM species separately in the presence of 1% seed of SO and MO using ThT binding kinetics (Figure 2E and 2F) respectively. Interestingly, we did not find any change in the aggregation kinetics of LMW species, specifically lag phase in the presence of 1% MO seed (Figure 2E), whereas, decrease of lag phase was observed in MM species aggregation kinetics in the presence of 1% MO seed. This data indicates 1% MO seed has no impact on LMW α-synuclein aggregation, on contrary, it recruit MM species and contribute to its aggregation. However, 1% SO seeds showed significant reduction of lag time in ThT kinetics indicating seeding assisted aggregation and impact was observed on MM species aggregation kinetics. Putting all results together, we conclude that glycation modulates the folding of α-synuclein and generates negatively charged species with greater exposed hydrophobic surface, leading to decrease in interaction with membrane and LMW species.

### Glycation affects internalization of α-synuclein species and modulates activation of microglia via its cytoskeletal structure

Microglia represents innate immune system of the brain that plays an important role in manages infection and inflammation (46). To understand the impact of conformationally different oligomers over internalization and subsequent, modulation of neuroinflammatory process in microglia, we treated microglia cells (BV2 cells) with 5 μM of different species of α-synuclein and measured the microglia activation. We stained the cells with IBA1 (red channel) and α- synuclein (green channel) after 12 h of post treatment with different α-synuclein species and analysed under SP8 confocal microscope (Figure 3). IBA1 is a cytoskeletal protein which is considered as marker of activated state of microglia (47, 48). Intensity of IBA1 (red channel) was found to be increased in case of MM treated cells, representing the activation of microglia cells. In case of LMW treated cells, IBA1 intensity was 10.68±1.4, whereas intensity was increased up to 12.64±2.2 upon treatment with MM (Figure 3F). Both MO and SO species were able to elevate the expression of IBA1, 13.0±2.0 and 13.0±1.7, respectively (Figure 3F). The expression of IBA1 was significantly high for both oligomeric species with respect to control BV2 cells and LMW treated cells. Upon interaction with any insult, microglia can rapidly activates and change its ramified morphology by reorganising its actin cytoskeleton, especially elongation of lamellipodia and filopodia (49). IBA1 is actin cross linking protein and previous research showcases its colocalization with filamentous actin towards lamellipodia in activated microglia (50). Therefore, to understand the effect of α-synuclein species over microglia morphology, we performed low force AFM with BV2 cells (Supplementary Figure 2A). We observed that microglia acquires morphology with expanded cytoplasm with increased processes, when treated with SO and MO species. Similarly, filamentous actin stained cells showed thickening of lamellipodia and increased branches upon treatment with SO and MO (Supplementary Figure 2B).

**Figure 3:**
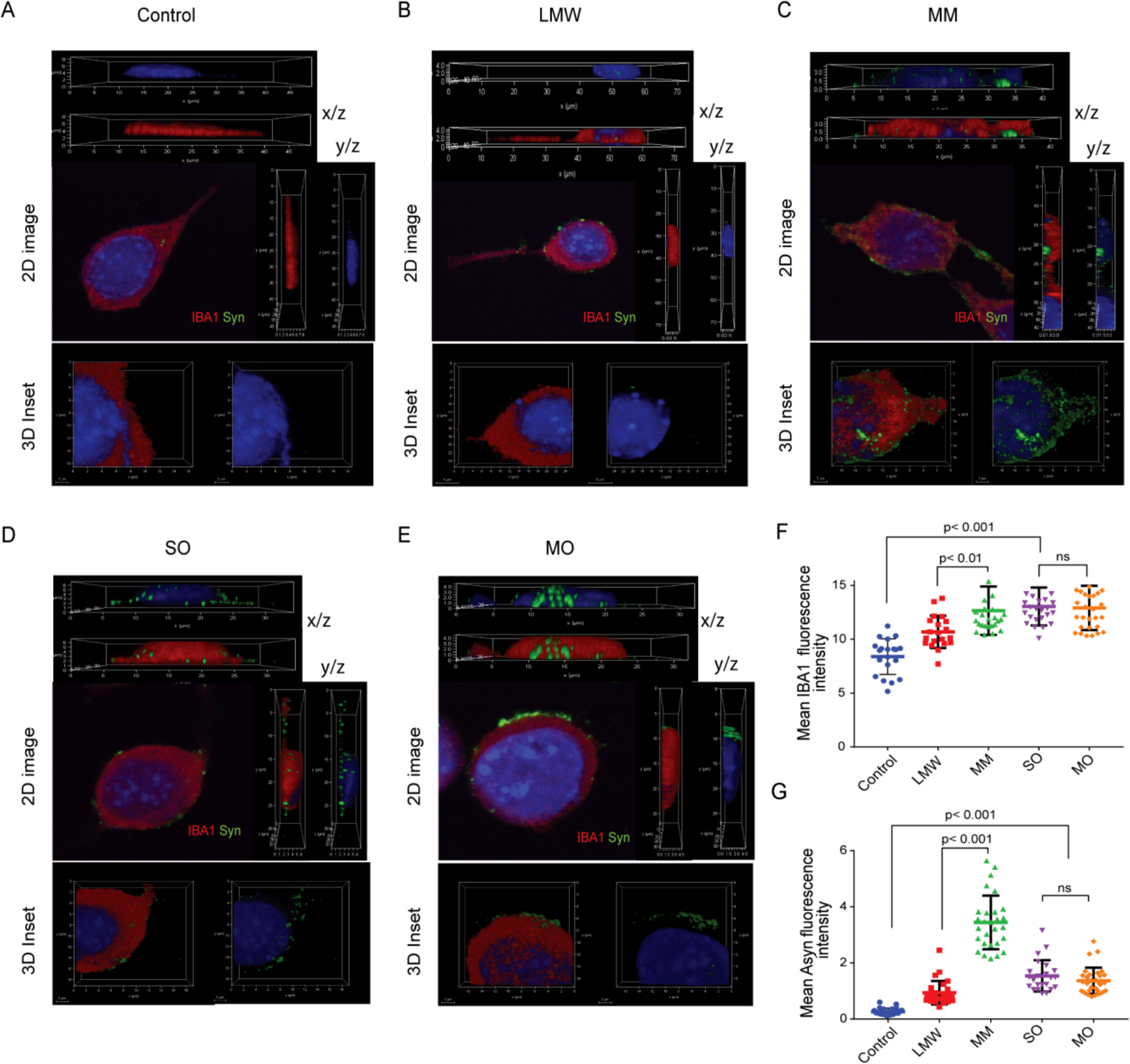
Glycation affects internalization of α-synuclein species and modulate microglia activation. (A-E) Confocal imaging representing 2D and 3D view of BV2 cells under treatment of LMW, MM, SO, MO α-synuclein species. Confocal images also supplemented with orthogonal view from different planes (x/y and y/z). BV2 cells stained with IBA1 (red channel) and α-synuclein (green channel). Scale bar, 10μm. (F) Quantitative analysis of IBA1 expression in cells under different treatments. (G) Quantitative analysis of α-synuclein expression in cells under different treatments, n>40 cells. Statistical significance was calculated using one way ANOVA followed by Tukey’s multiple comparison test in GraphPad Prism software.

In order to shed light on effect of glycation over internalization of these species inside the BV2 cells, we compare the intensity of synuclein species (green channel) in different z-stack, x/y plains and z-stacks based 3D constructed confocal images. We found that MM species were evenly distributed in the cytosol of BV2 cells than the LMW, which were clearly seen in 3D inset of Figure 3C and 3B, respectively. In case of oligomers treated cells, MO spices resided at the periphery of membrane whereas, more intensity was seen in cytosol in case of SO (Figure 3E and 3D, respectively). The intensity for α-synuclein species was calculated and represented in Figure 3G. We found that MM fluorescence intensity (3.44±0.95) was more than the LMW (0.93±0.42). We did no find much significance differences in overall fluorescence intensity between SO (1.53±0.55) and MO (1.36±0.46). With the help of mass spectrometry analysis, we were able to quantify the internalized synuclein species in BV2 cells. Our analysis indicates that internalized MM species were more than the LMW synuclein. However, SO were found to be internalized more than the MO (Supplementary Figure 3).

### Proteome analysis of microglial cells treated with glycated α-synuclein species shows unique protein expression pattern

Next we aim to understand the molecular changes in microglia upon exposure of α-synuclein species. We performed the quantitative proteomics analysis to address the microglial signalling mechanisms. We applied SWATH-MS to acquire the quantitative proteomics data from BV2 cells treated with LMW, MM, SO, MO. We acquired SWATH-MS data in three biological replicates and raw data were further analysed in Spectronaut pulsar. The molecular pathways and network analysis with the significantly regulated proteins were performed based on the data analysis workflow mentioned in Figure 4A. The raw data were normalized based on total ion count (TIC) of the SWATH-MS data (Figure 4B). Quality SWATH-MS data was assessed by coefficient of variance (CV), PCA plot, and Pearson’s correlation analysis. We found the median of coefficient of variance (CV) is < 14.7% (Figure 4C) and Pearson’s correlation value with > 0.85 with good reproducibility. All different treated cells were separated at the PC1 with a maximum 13.5% variation and PC2 with 27.3% variation, respectively (Figure 4D). PCA analysis of SO treated condition was separated from monomer treated condition and MO was shown to separate out from all other conditions.

**Figure 4:**
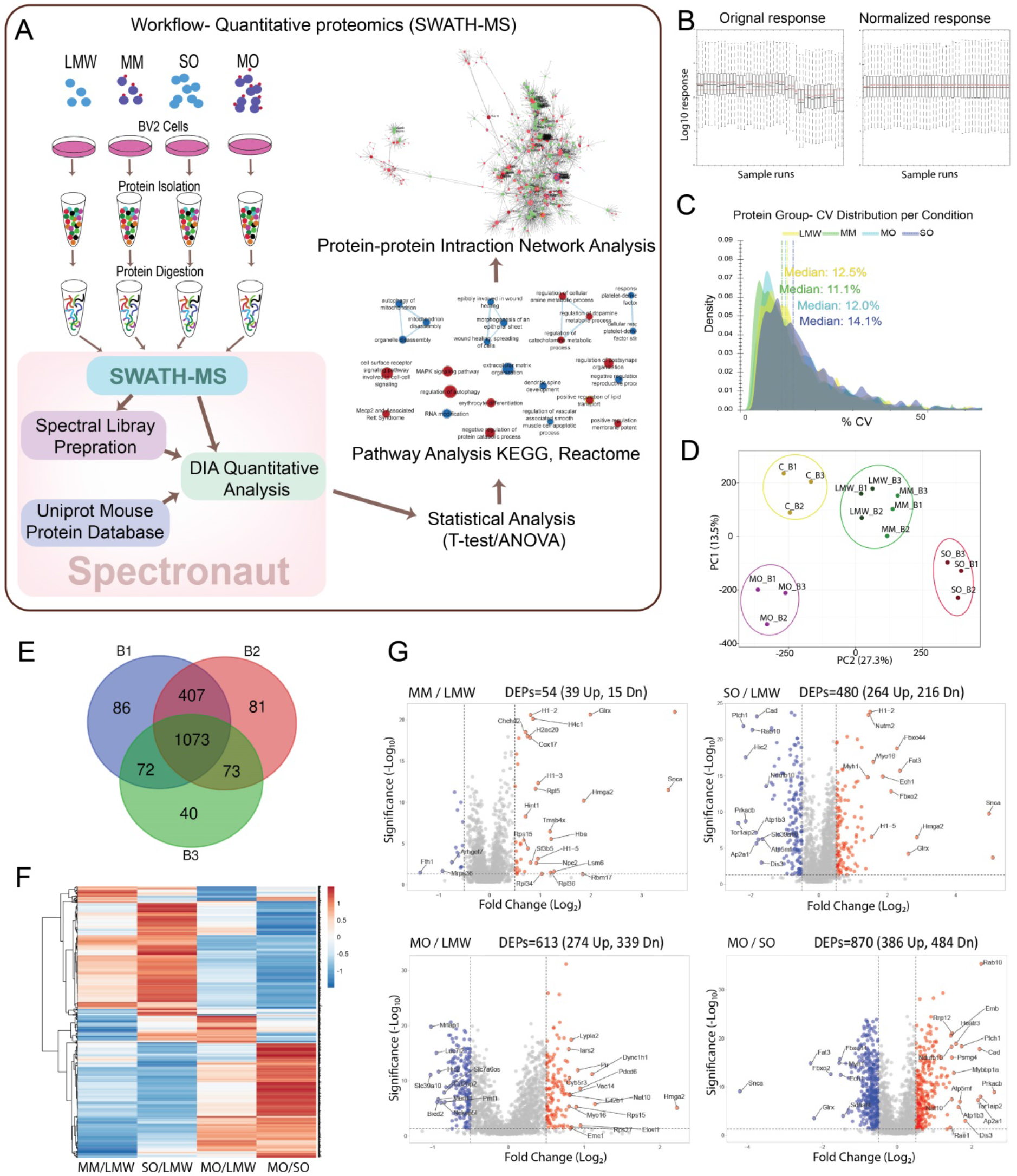
The quantitative proteomics analysis of α-synuclein species treated microglia cells. (A) Schematic representation of the workflow of SWATH-MS analysis. (B) Normalization of the SWATH-MS data based on Total Intensity Count (TIC) of MS/MS spectra. (C) Density plot showing the reproducibility of quantitative SWATH-MS data of replicates across treatment conditions, all treatment condition shows <15% coefficient of variation (CV) in the analysis. (D) Principal component analysis (PCA) of proteomics data with biological replicates indicated clear differences in proteomics profile of MO (pink circle) and SO (red circle) treated cell from monomer species (LMW and MM) treated cells (green circle). (E) Venn Diagram showing the common quantifiable proteins among biological runs. (F) Heatmap showing the relative expression profile of all proteins present in all biological samples. (G) Volcano plot representing the significant relative protein expression compared to LMW.

Data were further statistically analysed by applying t-test to compare between all α-synuclein species treated condition with LMW α-synuclein (LMW) treated condition. A total 1879 proteins were found to be significantly regulated in any pair of condition. As our primarily focus is to study the proteome modulation in MO versus SO condition, we observed a total 1073 proteins were significantly (p-value <0.05) changed in all biological replicates of MO/SO (Figure 4E). After applying expression cut-off (fold change ± >1.3), a total 870 proteins were found to be differentially modulated in MO/SO (Figure 4G). With respect to LMW treated cells, we found highest number of differentially expressed proteins in MO treated cells (MO/LMW, DEPs= 613) (Figure 4G). Heatmap analysis also revealed that the most of the proteins of SO/LMW oppositely regulated in MO/LMW (Figure 4F). Modulated proteins of MO/SO were further enriched with biological processes and pathway database.

### Gene ontology and pathway enrichment analysis reveals the unique regulation of specific biological pathways in MO/SO

All modulated 870 proteins of MO/SO were subjected to Gene set enrichment analysis (GSEA). We processed the data with Reactome mouse database pathway using all significant proteins (p-value <0.05) without any fold change cut-off filter and data were visualized in Cytoscape-Enrichment map with auto annotation to produce group enriched terms (Supplementary Figure 4). We found enriched pathway nodes of 413, 267, 98 and 392 for MM/LMW, SO/LMW, MO/LMW and MO/SO respectively. The enriched pathways were majorly linked to secretory pathway, actin dynamics, inflammation signalling events and Golgi related pathways based on the enrichment analysis of all significant proteins from all treated condition. MO/LMW showed unique pattern of upregulated enriched pathways in comparison to MM/LMW and SO/LMW. Interestingly, when we compared MO/SO, most of the pathways were up-regulated and they were shown to be down-regulated in MM/LMW and SO/LMW condition. Further we performed pathway enrichment for MO/SO condition with significantly regulated (p-value <0.05 and FC ±>1.3) proteins. Here, we applied g:Profiler analysis with ranked proteins and searched against KEGG and Reactome pathway mouse database and found that the mitochondrial apoptotic pathway, ribosome biogenesis, de novo folding, hedgehog signalling, innate immunity, clathrin-mediated endocytosis and programmed death apoptosis pathways were regulated (Figure 5). The comparison of MO vs SO highlighted many up- regulated pathways that were associated with Parkinson disease, neurodegeneration, ferroptosis, and metabolism of RNA, antigen processing, cellular response to hypoxia, DNA damage response and innate immunity. Similarly down-regulated pathways were linked with spliceosome, protein processing and presentation, cellular response to stress, programmed cell death and apoptosis. We further explored and validated a few specific pathways that were uniquely regulated in MO treatment condition.

**Figure 5:**
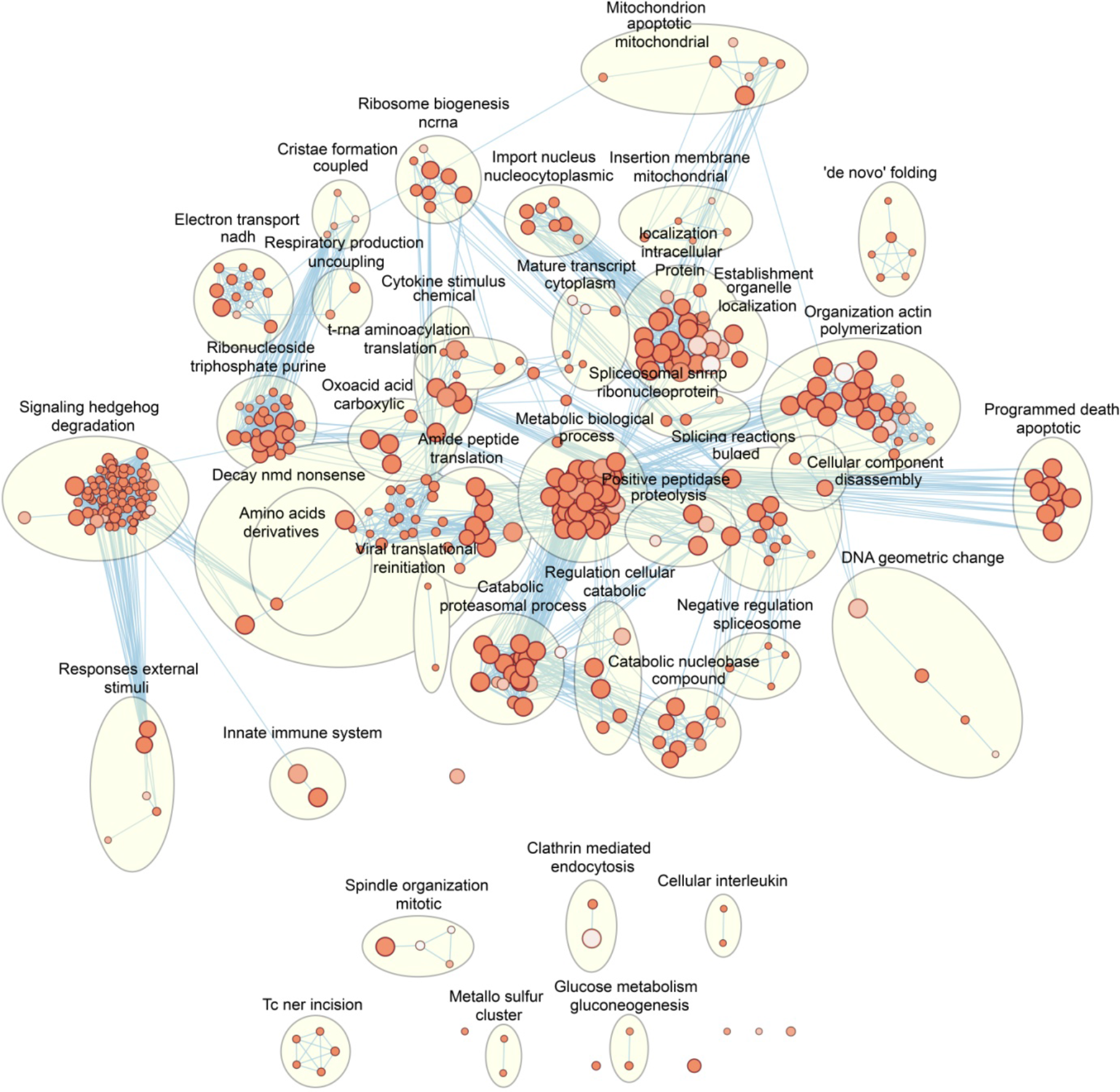
Pathway enrichment network of KEGG and Reactome based pathay database. All modulated proteins were used for pathway enrichment pathway enrichment for MO/SO condition using g:Profiler. The Enrichment Map shows the enriched gene-sets as a node. Edges between nodes overlap of member genes between pathway terms. AutoAnnotate function was applied to group gene-sets those belong similar category and pathways are depicted as circles.

### Glycated synuclein species prefer clathrin mediated cellular entry mechanism in BV2 cells

α-Synuclein and its different species enter the cells through different mechanisms. Passive diffusion through plasma membrane has been established in case of monomeric α-synuclein (51). However, α-synuclein species enter the cells mainly through endocytic pathways that include lipid raft mediated endocytosis, clathrin mediated endocytosis and phagocytosis (52, 53). Our quantitative proteomics data provided a strong support that endocytosis of α-synuclein species is a key mechanism in microglial cell entry. Similar to the previous reports, we also found an up-regulation of lipid-raft pathway related SERA protein in LMW treated condition. Interestingly, we observed a shift of internalization pathway of α-synuclein from lipid raft to clathrin dependent endocytosis due to glycation of synuclein. MM species showed up- regulation of clathrin dependent pathway. Recent report suggests that the internalization of α- synuclein to cells is conformation specific. So, it prompts to find the cellular entry mechanism of glycated oligomeric species. Here we closely viewed clathrin dependent endocytosis as this process was up-regulated in MO or SO condition compared to MM or L. We assessed the expression levels of dynamin, clathrin and AP2 proteins in our mass spectrometry dataset (Figure 6A). Dynamin, Clathrin and AP2 proteins mediate the interaction between clathrin and cargoes (54). Further investigation within the oligomeric species supported that the expression of dynamin and AP2 was found to be more up-regulated along with clathrin in MO treated condition (Figure 6A). Thus our mass spectrometry data provided a strong clue that clathrin mediated endocytosis is the preferred mechanism for glycated synuclein species.

**Figure 6:**
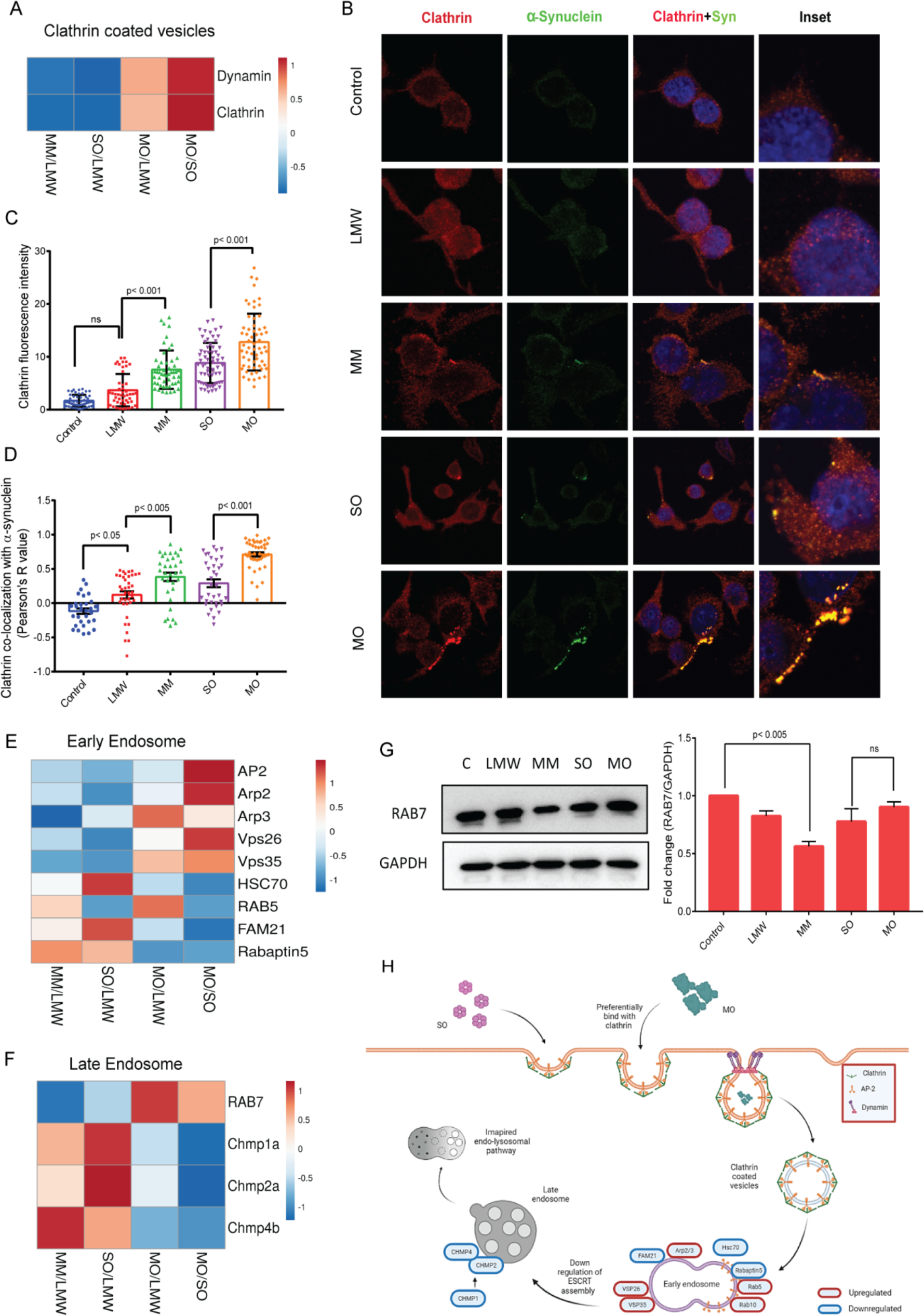
Glycation of α-synuclein modulate its endocytosis. (A) Heatmap representing the expression of clathrin coated molecules (Dynamin, clathrin and AP2) identified in mass spectrometry analysis. (B) Confocal microscopy images of α-synuclein species treated BV2 cells representing the expression of clathrin (Red channel) and α-synuclein (Green channel) as well as their colocalization. Scale bar, 5μm. (C and D) Quantitative analysis of clathrin intensity and its colocalization with α-synuclein in BV2 cells, n> 40 cells. Statistical significance was calculated using one way ANOVA followed by Tukey’s multiple comparison test in GraphPad Prism software. (E and F) Heatmap representing the expression of early and late endosome assembly molecules identified in MS analysis. (G) Western blot analsysis demonstrating the expression of RAB7 in α-synuclein species treated BV2 cells, n=3. Statistical significance was calculated using one way ANOVA followed by Tukey’s multiple comparison test in GraphPad Prism software. (H) Holistic representation of pathway observed in MO treated BV2 cells. Encircled red and blue molecules represent upregulated and downregulated protein, respectively.

In order to validate this observation, we treated the BV2 cells with different α-synuclein species, probed with clathrin antibody and observed under confocal microscope. We found an evidence of colocalization between α-synuclein and clathrin in both MO and SO treated cells (Figure 6B). However, the clustering of clathrin was seen significantly high on to the membrane in MO treated condition (Figure 6B and 6C). A highest colocalization α-synuclein (Pearson’s coefficient ∼0.9) (Figure 6D) with clathrin was observed in MO treated condition.

In canonical clathrin-dependent pathway, clathrin coated vesicles take up the cargoes to the lysosome for degradation via early and late endosome. Thus, we were interested in scouting the fate of MO species if these species were taken up for lysosomal degradation or any other pathways. We observed the up-regulation of early endosome regulator molecules (Arp2/3, VPS26, VPS35, Rab5 and rab10 in MO treated cells (Figure 6E) and down regulation of Hsc70, FAM21, rabaptin5 and these proteins are responsible for translocation of substrate to the lysosome for degradation (Figure 6E). Further, we analysed the expression of RAB7 which is responsible for maturation of early endosome to late endosome in our MS data and western blot. We found expression of RAB7 was significantly downregulated in MM treated microglia cells, however unchanged expression was observed in MO treated cells (Figure 19F and 6G). We also found downregulation of endosome sorting assembly i.e. ESCRT assembly in MO treated cells compared to SO treated cells. The endosomal sorting complex is necessary for transport (ESCRT) assembly which sorts the ubiquitinated proteins for lysosomal degradation (55). We found lowered expression of different charged multivesicular body protein (CHMPs) such as CHMP1a, CHMP2a, and CHMP4b in MO treated condition than the SO treated condition (Figure 6F). Collectively, our data provided evidence that the clathrin mediated endocytosed MO species were halted for its degradation and it may cause a proteostasis burden for the cells.

### Glycated α-synuclein oligomers alter mitochondrial function by promoting destabilization

Mitochondrial impairment is the pathological signature of Parkinson’s disease (56). Mass spectrometry analysis between MO and SO showed an up-regulation of mitochondrial respiratory chain complex 1, 2, and down regulation of complex 3, 4, 5 (Figure 7A). The down- regulation of complex 3 and 4 may be due to down-regulation of key molecules COX and Cytochrome-c (Figure 7A). Next, we analysed the expression levels of mitochondrial fission protein, Drp1 and mitochondrial fusion protein, Mfn2 in our mass spectrometry data (Figure 7B) and validated through western blots (Figure 7C). We found significantly high expression of Drp1 and moderate increase in Mfn2 expression under MO treatment condition. We further observed the upregulation of OMM proteins such as TOM22, TOM40 and downregulation of IMM proteins such as TIM10, TIM13, TIM8, TIM9 in MO compared to SO treated condition (Figure 7D and 7E). The fluctuation in mitochondrial transport machinery might alter the mitochondrial function, leads to its impairment. We also accessed the thioredoxin reductase 1 and thioredoxin level both oligomeric treatment conditions. Thioredoxin dependent system plays a crucial role in maintaining the cellular redox homeostasis (57). The expressions of thioredoxin reductase1 (TxnR1) and thioredoxin proteins (cytosolic (Txn) and mitochondrial (Txn2)) were found to be down regulated in MO than the SO treatment condition (Figure 7F). Importantly, mitochondrial HSP10/HSP60 the assembly, which requires for maintaining the levels of its client protein such as superoxide dismutase 2 (SOD2), was also found to be down regulated in MO treated condition (Figure 7F). Cumulatively, these results indicate that there is an increased mitochondrial oxidative stress upon treatment with MO species. To understand the resultant of this imbalance in the redox assembly, we measured the reactive oxygen species levels (ROS) in BV2 cells under different treatment conditions and we observed an increased cytosolic ROS in oligomers MO treated condition (Figure 7G). As reported, overproduction of ROS can activate the CyPD (Cyclophilin D) which leads to opening of mPTP (mitochondrial permeability transition pore) via ANT (ADP/ATP translocase 1) (58). In our MS data we observe the downregulation of the mPTP opening in MO treated condition. We found the upregulation of CyPD, ANT, BAX in MO condition (Figure 7H). We also found downregulation of Cyto C as well as the mPTP opening was downregulated in MO treated condition. We validated the MS observed expression of BAX and Cyto C via western blot (Figure 7I). Taken together, these results pointing out that the apoptotic pathway might come to rest in MO treated microglia cells.

**Figure 7:**
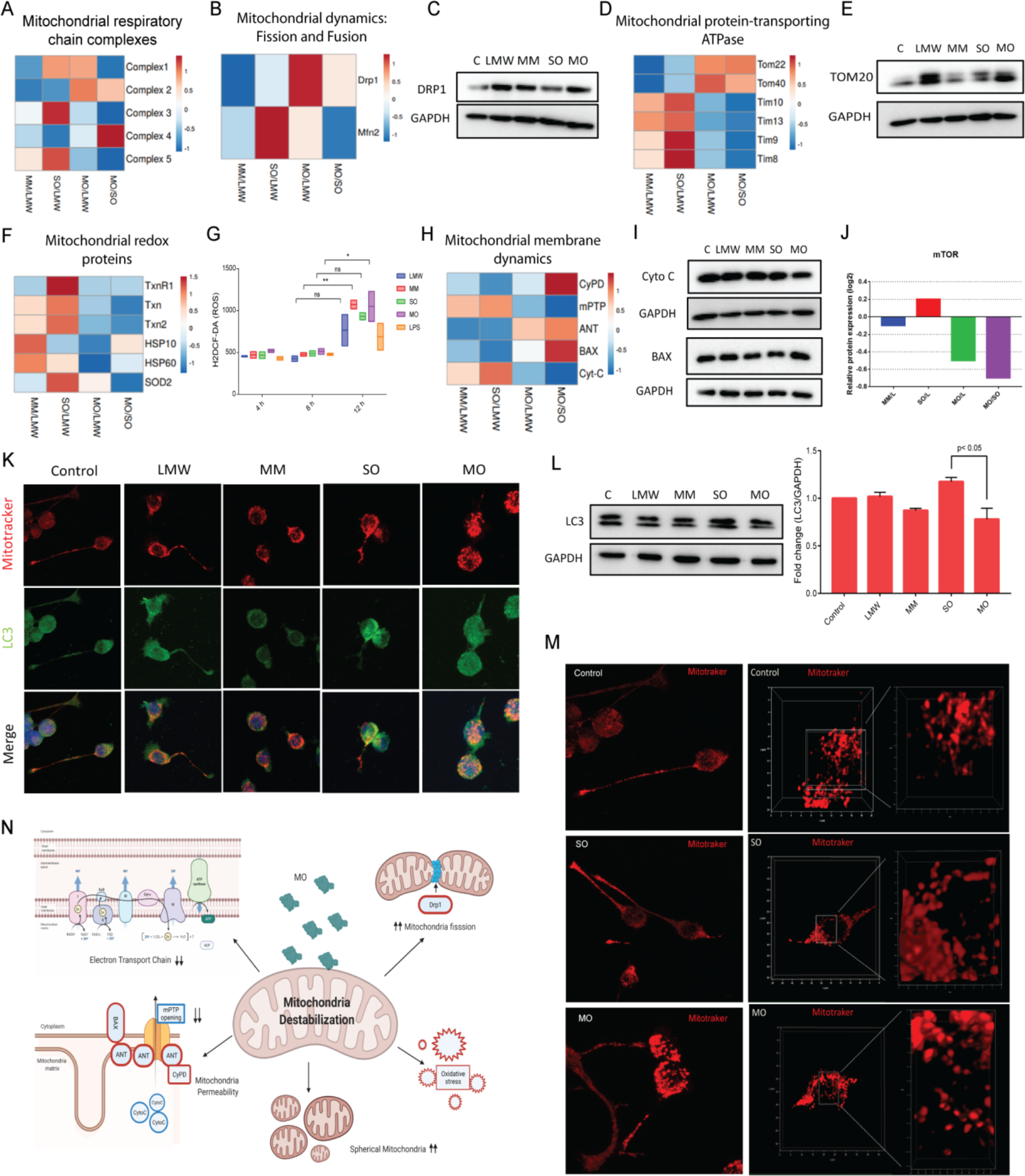
MO species promote mitochondrial destabilization. (A) Heatmap representing the expression of mitochondria respiratory chain complexes (Complex 1-5) identified in mass spectrometry analysis. (B) Heatmap representing the expression of mitochondria dynamics protein (Drp1 and Mfn2) identified in mass spectrometry analysis. (C) Western blot showing the expression of Drp1 and Mfn 2 in BV2 cells under different treatment conditions. (D) Heatmap representing the expression of mitochondria protein-transporting ATPase (Tom 22, 40 and TIM10, 13, 8, 9) identified in mass spectrometry analysis. (E) Representative western blot demonstrating the expression of TOM20 in BV2 cells treated with α-synuclein species. (F) Heatmap representing the expression of mitochondria redox related proteins (TxnR1, Txn1, Txn2, HSP10, HSP60, SOD2) observed in MS data. (G) H2DCF-DA staining of BV2 cells demonstrating the generation of ROS (reactive oxygen species) upon treatment of α-synuclein species. (H) Heatmap representing the expression of mitochondria membrane dynamic related proteins (CyPD, mPTP, ANT, BAX, Cyto C) observed in MS data. (I) Representative western blot demonstrating the expression of BAX and Cyto C in BV2 cells treated with α-synuclein species. (J) Graph representing the relative expression of mTOR quantified in mass spectrometry. (K) Confocal microscopy images of α-synuclein species treated BV2 cells representing the expression of Mitotracker (Red channel) and LC3 (Green channel) as well as their colocalization. Scale bar, 10μm. (L) Western blot demonstrating the expression of LC3 in BV2 cells treated with α-synuclein species. Quantification of LC3 protein levels in BV2 cells demonstrate significant increase in LC3 in SO treated BV2 cells. Statistical significance was calculated using one way ANOVA followed by Tukey’s multiple comparison test in GraphPad Prism software. (M) Confocal microscopy images of α-synuclein species treated BV2 cells representing the mitochondria stained with mitotracker (Red channel). 3D construction of mitotracker stained cells representing the shape of mitochondria in SO and MO treated cells, n> 25 cells. (N) Graphical representation of mitochondrial pathway observed in MO treated BV2 cells. Encircled red molecules represent upregulated protein.

Next, we were interested to know the fate of altered mitochondria. ANT is a positive regulator of mitophagy (59) and it was upregulated in MO treatment condition (Figure 7H). We also found downregulation of mTOR, which controls autophagy and mitophagy (Figure 7J). Further, we stained the cells with mitotraker (Red channel) and LC3 antibody (Green channel) upon α-synuclein species treatment and viewed under confocal microscope (Figure 7K). LC3 is responsible for formation of autophagosome membrane (60). However, LC3 expression was significantly higher and localized with mitotracker in SO treated cells than MO treated cells (Figure 7L and 7K). Next, we did the 3D construction of mitotracker stained mitochondria in confocal microscopy and observed change in mitochondria shape from elongated to round form in MO treated cells (Figure 7M). Collectively, our results indicate that MO cause more mitochondrial destabilization than the SO and mark less it for mitophagy.

### Glycated α-synuclein oligomers activate the microglia cells preferentially via Inflammasome pathway

Recent studies have shown that extracellular α-synuclein aggregated species activate microglia cells via pattern recognition receptors i.e toll-like receptors (TLRs) (61). Specifically, oligomeric forms of synuclein are reported to interact with TLR2 and TLR4 on the surface of glia cells and activate the TLR signalling (33, 36, 62). First, we tested if glycated α-synuclein and its oligomeric species can bind to TLR2 and activate the TLR2 pathway. We checked for *in-vitro* binding of glycated species of synuclein with TLR2 using BLI. TLR2 were loaded over AR2G sensor tips and incubated with species of α-synuclein. Interestingly, glycated α-synuclein hinders TLR2 binding both in monomeric as well as oligomeric form (Figure 8A). The Kd values for respective binding graphs are represented in **Supplementary Table 3**. TLR2 showed good binding with LMW and SO species showed strongest binding with KD= 1.50E- 07±2.5 M. We found a substantial loss of TLR2 binding on glycation of α-synuclein in both MM as well as MO species. Here we used PAM3CSK4 as a positive control and it showed a strong binding with KD of 1.67E-06±2.2 M. To enhance our understanding, we used HEK- Blue™-hTLR2 cells which harbour SEAP reporter gene under the control of promoter fused to NF-κB. The level of SEAP is a direct representation of production of NF-κB. We treated the cells with synuclein species and monitored the level of SEAP by measuring its absorbance at 620 nm in real time. We found the SEAP levels were highest in PAM3CSK, a well-known agonist of TLR2 (Figure 8B). The LMW species showed basal level activation of NF-κB. However, SO treated cells showed highest TLR2 activation and it was almost 3 times higher than LMW specie. On the other hand, both with MM and MO species showed significant reduced activation of NF-κB compared to SO. Though both MM and MO species showed 1.5 fold higher NF-κB activation than LMW species. However, it is unclear despite impaired binding in BLI how glycated Monomer and Oligomers shows considerable activation TLR2 in HEK-Blue™-hTLR2 cells. Both in-vitro TLR2 binding experiments using BLI and TLR2 activation in HEK-Blue™-hTLR2 cells suggest that glycation of synuclein reduce it binding to TLR2, simultaneously reduction in activation of Nf-κB pathway.

**Figure 8:**
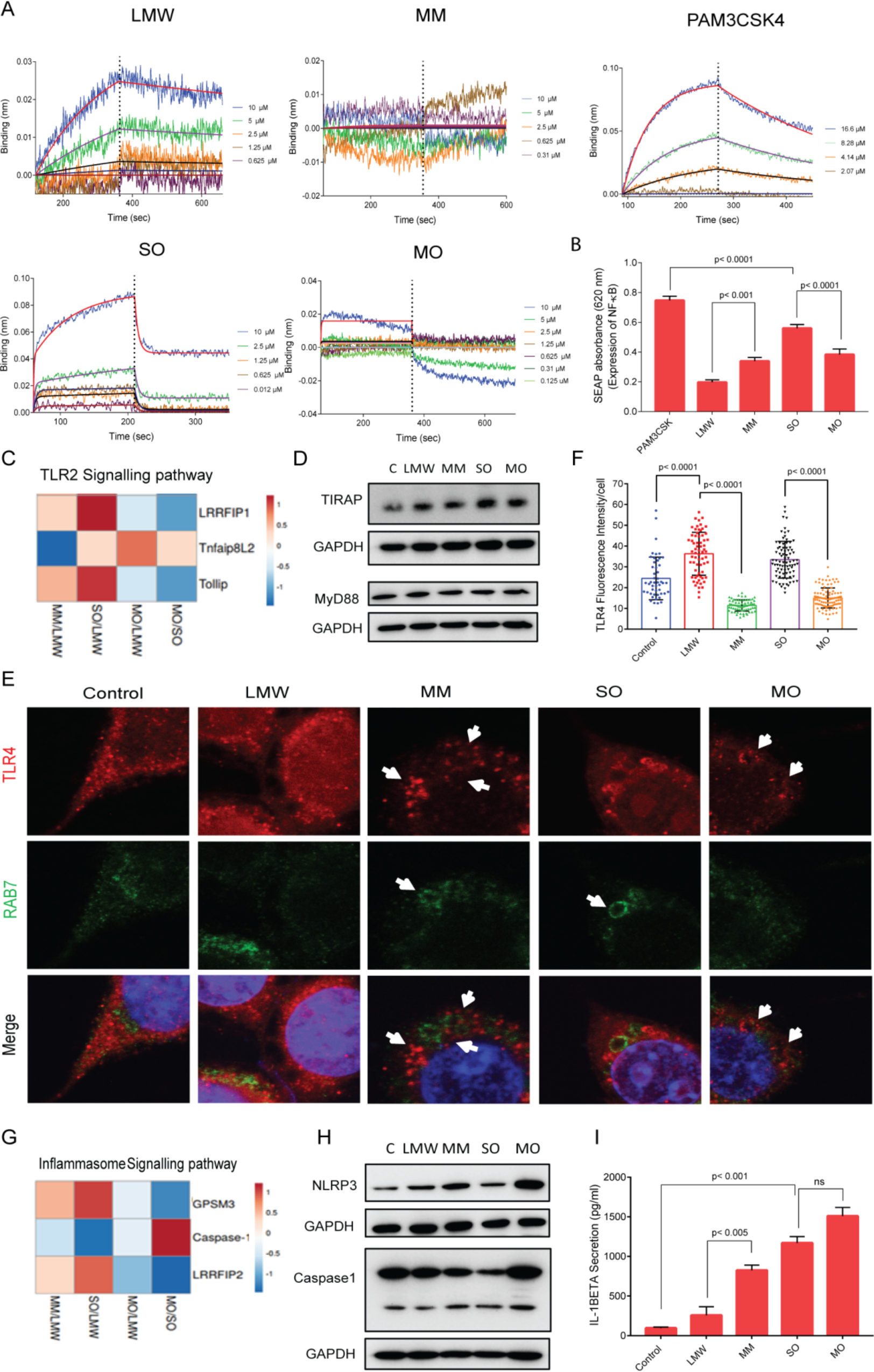
Glycated species modulate the inflammatory response of microglia. (A) BLI graph showing the interaction of α-synuclein species with TLR2 by loading TLR2 purified protein over AR2G sensor tips and dipped in wells containing 10 μM to 0 μM of α-synuclein species. Among all species, LMW, SO and PAM3CSK4 (known agonist of TLR2) interacted significantly more to AR2G sensor loaded with TLR2. (B) HEK-Blue™-hTLR2 cells, harbouring SEAP reporter gene under the control of promoter fused to NF-κB, treated with α-synuclein species. The levels of SEAP were monitored by measuring its absorbance at 620 nm. SEAP levels were higher in SO and PAM3CSK4 treated cells, whereas, significantly lesser expression were seen in MM and MO treated cells. Statistical significance was calculated using one way ANOVA followed by Tukey’s multiple comparison test in GraphPad Prism software. (C) Heatmap representing the expression of TLR2 signalling regulating proteins (LRRFIP1, Tnfaip8L2 and tollip) observed in MS data. (D) Western blot demonstrating the expression of TIRAP and MyD88 in BV2 cells treated with α-synuclein species. TIRAP protein levels in BV2 cells demonstrate significant increase in SO treated cells; however, MyD88 expression was unchanged in all treatment conditions. (E) Confocal microscopy images of α-synuclein species treated BV2 cells representing the expression of TLR4 (Red channel) and RAB7 (Green channel) as well as their colocalization. Scale bar, 5μm. (F) Quantitative analysis of TLR4 intensity in BV2 cells treated with α-synuclein species, n> 50 cells. Statistical significance was calculated using one way ANOVA followed by Tukey’s multiple comparison test in GraphPad Prism software. (G) Heatmap representing the expression of regulating proteins of NLRP3 inflammasome signalling (GPSM3, Caspase1 and LRRFIP2) observed in MS data. (H) Western blot demonstrating the expression of NLRP3 and Caspase1 in BV2 cells treated with α-synuclein species. NLRP3 and caspase1 protein levels in BV2 cells demonstrate significant increase in MO treated cells. (I) ELISA of IL-1β representing levels of secreted IL-1β by α-synuclein treated BV2 cells, n=3. Statistical significance was calculated using one way ANOVA followed by Tukey’s multiple comparison test in GraphPad Prism software.

To get better insight in neuroinflammation process, we checked the quantitative mass spectrometry data. We found upregulation of certain TLR2 pathway molecules like LRRFIP1 and TOLLIP in SO treated condition compared to LMW (Figure 8C). Both these molecules act as a regulator of NF-κB. Interestingly, downregulation of these molecules were observed in MO treated condition in comparison to LMW. We further verified both expression of TLR2 signalling downstream molecules including like myeloid differentiation primary response gene 88 (MyD88), TIR domain containing adaptor protein (TIRAP) (Figure 8D). BV2 cells were treated with LMW, MM, SO and MO species for 6 h and probed the expression of these molecules in western blot. We used 6h of treatment to probe the early regulatory molecule of TLR activation pathway via western blot analysis. We detected slight increase of TIRAP expression upon exposure of SO species in comparison to other treated conditions. However, MyD88 protein expression was unchanged in all cases (Figure 8D). Collectively, these results confirm that SO species prefer to activate TLR2 mediated pathways, however, MO species showed lesser preference to TLR pathway. In addition, we also checked the effect of glycated species with respect to TLR4 mediated inflammation. We treated BV2 cells with α-synuclein species and probed for expression of TLR4 via confocal microscopy. Interestingly, we observed internalization of TLR4 in MM and MO treated cells (Figure 8E). It was reported that endocytic pathways, more precisely dynamin and clathrin-mediated regulate the TLR4 signalling (63). TLR4 localization on early/late endosome was documented to regulate the TLR4 mediated adaptive immunity (64). So, we checked TLR4 localization with RAB7, a late endosome marker. We observed the arrangement of TLR4 protein in circular shape and it was co-localized with RAB7 consisting endosomes. Subsequently, we quantified the intensity of TLR4 in all treatment conditions and found significant increase in TLR4 expression in LMW and SO treated BV2 cells in comparison to untreated cells. Interestingly, its expression was significantly reduced in MM and MO treated cells (Figure 8F). We hypothesise that MM and MO species increase TLR4 endocytosis via clathrin dependent pathway and target it for lysosome degradation.

Previous reports explicate that the dysfunction of lysosome machinery and mitochondrial degradation can activate NLRP3 inflammasome (65). ROS and Ca^2+^ signalling also have critical role in activation of NLRP3 inflammasome (66, 67). This made us curious to know whether mitochondrial destabilization by MO species can shift TLR mediated inflammatory pathway to NLRP3 inflammasome mediated pathway. Interestingly we observed an upregulation of NLRP3 inflammasome pathway molecules like caspase1 levels were observed in MO treated condition in comparison to SO (Figure 8G). The negative regulators of NLRP3 such as and LRRFIP2, HSPA8 and GPSM3 were down regulated (Figure 8G, Supplementary Figure 5) in MO vs SO condition. To verify the NLRP3-inflammasome mediated neuroinflammation, we treated BV2 cells with α-synuclein species for 6h and probe for the expression of NLRP3 and caspase1 via western blotting. We found increased expression of NLRP3 in MM and MO species treated cells (Figure 8H). The expression and cleavage of caspase1 were higher in MO treated condition. Further, we measured the secreted IL-1β levels using ELISA. We observed highest IL-1β secretion in MO treated condition. We found 2.0, 2.5 and 3.5 folds increase of IL-1β secretion of MM, SO and MO species treated condition respectively in comparison to LMW species treated condition. However, secreted IL-1β levels were not changed significantly in MO species with respect to SO treated cells (Figure 8I).

Taken together, our data strongly suggest that glycation of α-synuclein species reduces it binding to TLR2, compromises activation of Nf-κB pathway. In the same time glycated α- synuclein species potentiate the neuroinflammation preferentially by activating the NLRP3 inflammasome pathway.

## Discussion

Parkinson’s disease is often characterized by misfolding and aggregation of α-synuclein, followed by degeneration of dopaminergic neurons (2). Significance efforts have been made towards understanding the fibrillar and oligomeric species that accumulate within the cell and cause neuronal toxicity (11, 68). Strong evidences support the abundance of α-synuclein oligomeric species in-vivo, however, structural characterization of toxic oligomers is challenging because of their transient nature and size variability (68, 69). α-Synuclein strains are different in structure, level of toxicity, seeding and propagation properties (70, 71). Such strains may account for difference in disease propagation in different individuals with other synucleinopathies. Here, our study uncovered the impact of clinically identified PTM i.e glycation over the morphology, aggregation and membrane binding of α-synuclein. We have shown that mainly N-terminal and post-NAC domain lysine residues are modified by methylglyoxal and this is consistent with earlier observation. In synergy to previous reports, we have also shown that methylglyoxal drives glycation and restricts α-synuclein in the oligomeric during aggregation conditions (23, 24). Despite of similar disordered structure of glycated LMW α-synuclein, glycated oligomers exhibit lower beta rich structure than unmodified oligomers. Both AFM and TEM analysis of MO and SO species also provide an evidence of deviations in its ultrastructure. SO species shows donut shaped oligomers. However, MO do not show regular geometrical rearrangement and they are comprised of sandwich 2-4 units. Similarly surface topology and charges of MO also significantly differ from SO. Glycated α-synuclein species show higher exposure of hydrophobic surface and carries significantly high negative charge at the solvent surface interface. Both solvent exposed hydrophobic surface and net surface charges can directly influence α-synuclein aggregation kinetics and its membrane binding. LMW species show intense interaction with the lipid membrane through their N-terminal positive charge lysine residues. The MGO modification of N-terminal lysine residues impose a negative charge on these lysine residues and prevent membrane binding. These negative charges interferes phospholipids head group interaction. Fibrillation of any amyloid proteins is a multistep process, precedes through initial nucleation which further recruit bulk unit and generates protofibrils and subsequent fibrils (41, 43, 45). The seed amplification is the important mechanism in amyloid fibril structure formation (43). Thus, glycation of α-synuclein may not hamper the nucleation but certainly inhibits amplification step. This may be due to surface charge compatibility between seed and LMW species.

Recently, it is reported that microglia prevent neurodegeneration by clearing neurone released synuclein species via selective autophagy (30). We have attempted to delineate the mechanism through microglial respond towards different wild type and glycated species. First, we checked cellular entry mechanism. We find SO species entre the microglia cells more efficiently. However, MO species appear to be predominantly localized at the plasma membrane. Despite the differential cellular entry, both SO and MO species activate the microglia (based on IBA1 expression) in similar extent. With the help of quantitative proteomics, we identify that the influx of MO in microglia cells is predominantly mediated through clathrin-dependent endocytosis and colocalization of clathrin are seen with MO. It is well documented that clathrin coated vesicles fuse and transfer their cargos to Rab5 positive early endosomes (54, 72). Thereafter maturation of early endosome into late multivesicular endosome, cargoes is finally sorted by the ESCRTs machinery for their lysosomal degradation (55). Upregulation of the ESCRT assembly in SO treated microglia cells suggest its intracellular trafficking for degradation. In contrast to SO, MO species somehow escape the process of endosome trafficking as ESCRT assembly was downregulated. In neuronal cells, it has been reported that MGO treatment cause failure of proteasome assembly and contribute to α-synuclein accumulation (23). Here, we can conclude our findings that glycation of α-synuclein may cause defects in endo-lysosomal pathway, hence, reduced clearance can promote accumulation of MO in microglia.

α-Synuclein is reported to interact with mitochondrial membrane proteins such as Drp1 and TOMs and cause mitochondrial destabilization (73, 74). Our findings prove that MO hampers the ATP generation machinery and destabilise the mitochondria membrane potential. Simultaneously, it also increases the mitochondrial fission process by increasing the expression of Drp1 and also generates the mitochondrial redox stress by altering the redox homeostasis. Another molecule such as Bax (a Bcl-2 family protein) which facilitate the opening of mPTP, can cause the cave in of the mitochondrial membrane potential and release of pro-apoptotic mediators takes place such as cytochrome c (56). MO showed lesser release of cytochrome c due to downregulation of mPTP, indicating halting of apoptosis process. Mitophagy is a mitochondrial homeostatic mechanism by which it clears the damaged mitochondria (75). However, upregulation of mitophagy were seen in SO treated cells as autophagy marker, LC3 were more colocalizated with mitotracker than microglia cells treated MO.

Activation of microglia cells and pro-inflammatory cytokines release are the key factors that contribute in pathophysiology of Parkinson’s disease (27-29). α-Synuclein known to provoke inflammatory response via toll-like receptors (TLRs) especially TLR2 and TLR4 (32, 33, 61). We first revisited the TLR mediated inflammatory response by α-synuclein by using TLR2- expressing HEK cell lines. In synergy to previous reports, SO species increases expression of NF-κβ, an downstream molecules of TLR2 pathway. Whereas, significant decrease in NF-κβ expression was observed in MO species treated cells. To our surprise, TLR2 interaction studies also reveal that MO and MM have weak binding towards TLR2 than SO and LMW, respectively. Moreover, we observed downregulation of TLR2 regulatory molecules in MO treated cells in comparison to SO species. Our study uncovered a distinct feature of MM and MO species triggered immunity is that they enhances TLR4 endocytosis and subsequently, probe it for degradation. Endocytosis of TLR4 is important for regulation of adaptive immunity (64). So far, LPS and ethanol are known to trigger TLR4 endocytosis and activate the NF-κβ and MAPK pathway, whereas it is not observed with α-synuclein treatment (30, 63). Future studies should be conducted to conclude the significance of glycated α-synuclein species mediated TLR4 endocytosis. However, we can speculate based on available literature and our observations that it is regulated by clathrin-dependent endocytic pathway (63). Next, we looked into another aspect of immunity i.e NLRP3-inflammasome dependent. α-Synuclein can stimulate the NLRP3 inflammasome, which promote caspase-1 activation, followed by maturation and release of IL-1β and IL-18 pro-inflammatory cytokines (77, 78). Our results showed strong preferences MO species for the activation of NLRP3-inflammasome pathway, as increased expression of NLPR3 and caspase1 were observed upon MO treatment. Activation of NLRP3-inflammasome machinery is reported to prime by TLR2 and TLR4 dependent NF- κβ release (79). Collectively, we observed lesser expression of NF-κβ and higher activation of NLRP3-inflammasome pathway at early time point of treatment. With this we can conclude that NLRP3-inflammasome activation is independent of TLR-NF-κβ priming. Future studies can be conducted to shed light on the glycated species driven NLRP3-inflammasome mechanism. Our data strongly suggest that MO conformation has significant role in dictating the inflammatory pathway in microglia.

Taken all together, we can summarize that glycation of α-synuclein drive the microglia response differently than the synuclein. Our observations have also raised some open questions which can be address in future to understand the underling mechanism of Parkinson’s disease. First, how glycation of α-synuclein dictate the arrangement of monomers and form oligomers without compromising with its secondary structures. Second, what is the underlying mechanism for escape of glycated synuclein oligomers from endo-lysosomal pathway can be another aspect which provides deeper understanding of PD.

## Conclusions

Through our study we addressed a persisting problem of the field, which is, α-synuclein conformation specific microglial inflammatory response. Our results provide detailed morphological signatures of wild type α-synuclein and glycated α-synuclein oligomeric species. Glycated α-synuclein oligomers internalized into microglia cells via clathrin-dependent endocytosis, contribute to redox stress and impairment of mitochondrial dynamics. Glycated α-synuclein oligomers activate microglia preferentially via NLRP3-inflammasome pathway. Taken together, our study provided mechanistic feedback of microglia towards glycated α-synuclein oligomers. Further studies can be conducted to elucidate the role of glycated α-synuclein oligomers as a player of Parkinson’s disease pathology.

## Methods and materials

### Purification of α-synuclein and generation of glycated α-synuclein

Wild type α-synuclein plasmid 36046 (pT7-7) was generously gifted by Professor Hilal Lashuel [40]. It was further expressed in BL21 (DE3) strain of E.coli. Protein expression was induced using 1mM IPTG and protein purification was performed using ammonium sulphate precipitation method, very well established protocol for α-synuclein purification [41]. For long term storage, purified protein was lyophilized and stored at -80 ^0^C.

Low molecular weight species (LMW) of α-synuclein were prepared according to established protocol [42]. Lyophilized α-synuclein was solubilized in glycine-NaOH buffer, pH adjusted to 7.4 using NaOH. Solubilised protein were passed through 100 kDa Amicon® cut-off filter and flow through collected for protein estimation. α-synuclein oligomer SO were prepared by incubating 200 μM of LMW in 10 mM phosphate buffer pH 7.4 at 37 ͦ C under agitation condition for 16h. For the preparation of glycated α-synuclein LMW (MM), α-syn LMW (200 μM) were incubated with 5 mM of methylglyoxal (MGO) solution in 10 mM phosphate buffer at 37 ͦ C under static condition for 20 h. Excess of MGO were removed using 10kDa Amicon® cut-off filter. Protein concentration was estimated and MM was characterized using mass spectrometry (MALDI). Glycated α-synuclein oligomers (MO) was prepared by incubating 200 μM of MM in 10 mM phosphate buffer pH 7.4 at 37 ͦ C under agitation for 24 h. Final oligomers concentration was determined by measuring its absorbance at 280 nm (extinction coefficient 5960 M^-1^ cm^-1^). Most of the experiments were performed with freshly prepared α- synuclein species. If required, species can be kept at -20 for 2-3 weeks, centrifuged at high speed to remove higher aggregates before experiment.

### Thioflavin T and ANS fluorescence

The aggregation kinetics of LMW and MM was monitored by ThT fluorescence in 10 mm x10 mm pathlength cuvette, using a HITACHI F7000 fluorescence spectrophotometer. Both species (200 μM) in 10 mM phosphate buffer (pH 7.4) with 0.1% sodium azide were incubated at 37 ͦ C with 180 rpm agitation. During the course of aggregation kinetic, 7 μM protein aliquots were collected at different time intervals, followed by incubated with 8 μM of ThT dye. ThT fluorescence of each sample was measured by exciting at 442 nm and emission spectrum recorded between 460 to 600 nm (5 nm slit widths). For secondary nucleation assay, ThT aggregation kinetic of LMW (50 μM) was set up in presence of 1% SO and MO seed with equimolar concentration of ThT in 96 well plate. ThT fluorescence was monitored at 30 min intervals in Tecan Infinite F200 Pro plate reader upon excitation at 448 nm and emission at 482 nm) . For ANS binding assay, each species of α-synuclein was incubated with ANS in 10 mM phosphate buffer pH 7.4 for 15 min. ANS fluorescence spectra was monitored by exciting at 350 nm and recording the emission spectrum between 400 nm to 600 nm.

### CD spectroscopy

Far-UV CD spectrum of α-synuclein species (5 μM) was monitored between 190-260 nm using 0.2 mm pathlength cuvette in JASCO J815 CD spectrophotometer. Six accumulations were taken for each sample and background subtraction was done using a 10 mM phosphate buffer pH 7.4. The final spectra were represented in molar ellipticity. Protein secondary structure was analyzed using DichroWeb, an online analysis tool for protein CD spectra. CDSSTR method with a reference set 7 was applied for data analysis.

### TEM imaging

All α-synuclein species were diluted in 10 mM phosphate buffer pH 7.4 at concentration of 15 μM. 5 μl of each sample were placed on a glow discharged carbon coated copper grid for 1 min. The extra sample from grid was blotted off and stained with uranyl acetate for 2 min. Stained grid washed thrice with milliQ water. Air dried sample grids were imaged in JEM1400 Flash (JEOL, Japan) transmission electron microscope at different magnifications.

### Atomic force microscopy of species of α-synuclein

For the preparation of slides for atomic force microscopy, samples were diluted in filtered milliQ at concentration of 10 μM. 20 μl of species of α-synuclein were applied on freshly cleaved mica and subsequently, excess of sample were washed with MilliQ. Under intermittent contact mode, Air dried samples were imaged by silicon cantilever (spring constant of 13-77 N/m), in JPK Nano Wizard III atomic force microscope. Cantilever had drive frequency of 300-320 kHz and the scan rate for image kept between 0.8-1 Hz. JPK processing software utilized for topographic AFM image processing.

### Preparation of Lipid Vesicles

DOPC(1,2-dioleoyl-sn-glycero-3-phosphocholine),DOPE(1,2-dioleoyl-sn-glycero-3- phosphoethanolamine) and DOPS (1,2-dioleoyl-sn-glycero-3-phospho-L-serine) lipid vesicles with mass ratio of 50:20:30 were prepared via sonication and extrusion method. The 200 μM of lipid mixture was dissolved in Chloroform: Methanol (2:1, v/v). The solvent was evaporated by purging a stream of N2 gas. The thin layer of lipid was then kept under reduced pressure for 8 h and then, hydrated with 10 mM phosphate buffer pH 7.4. The lipid vesicles were generated by 21 freeze/thawing cycles and 12 extrusion cycles through a 50 nm polycarbonate filter using Avanti® extruder. The size of vesicles was monitored by dynamic light scattering with a Zetasizer Nano ZS (Malvern Instruments).

### Biolayer interferometry (BLI)

BLI experiments were performed on an Octet RED Instrument (fortéBIO). Ligand immobilization, binding reactions, regeneration, and washes were conducted in wells of black polypropylene 96-well microplates. For interaction study between α-synuclein species and lipid vesicles, 20 μg of LMW and MM species were independently immobilized on amine-reactive biosensors (AR2G biosensors) in 10 mM acetate pH 5 buffer, using 1-ethyl-3-(3- dimethylaminopropyl)-carbodiimide (EDC) and N-hydroxysuccinimide (NHS). Equal concentration of lipids in 10 mM phosphate buffer pH 7.4 was used for binding with LMW and MM. Binding was carried out in 10 mM phosphate buffer pH 7.4 at 25 ͦ C with 120 sec association and 120 sec dissociation. Data were analyzed using Data Analysis (fortéBIO), with Savitzky-Golay filtering. The binding was fitted to 1:1 ligand binding model, the steady-state analysis was performed to obtain the binding kinetics contents (Kd). Under similar experimental conditions, binding of α-synuclein species with LMW were monitor and analysed. Concentration used for all α-synuclein species was 10 μM and binding curve were was fitted to 1:1 ligand binding model.

TLR2 binding with α-synuclein species were performed by immobilizing TLR2 on AR2G biosensors in 10 mM acetate pH 5 buffer, using amine coupling. Binding analysis of α- synuclein species (10 μM- 0 μM) was carried out at 25 ͦ C, 1000 rpm in 10 mM phosphate buffer (pH 7.4) containing 150 mM NaCl, with a 120 sec of association followed by a 120 sec of dissociation. The surface was thoroughly washed with the running buffer followed by dipping in regeneration solution. Data were analyzed and binding curve was globally fitted to a 2:1 heterogeneous and 1:1 ligand binding model, the steady-state analysis was performed to obtain the binding kinetics contents (Kd).

### Dynamic light scattering (DLS) and Zeta potential

DLS and zeta potential measurements of species of α-synuclein were performed using Zetasizer Nano ZS (Malvern Instruments). All α-synuclein species were spun at 13000 rpm for 30 min and quantified before DLS and zeta potential measurements. Size distribution analysis of all samples (10 μM) was done in 3 mm pathlength quartz cuvette at room temperature. Intensity- size distribution data were acquired and average of 4 independent measurements was reported. All samples were diluted upto 25 μM in 10 mM phosphate buffer and Zeta potential measured using DTS1060 electrode cuvette.

### Mass spectrometry

#### Trypsin digestion of MGO modified α-synuclein

For site identification, we performed in-gel digestion with trypsin and Glu-C enzyme. So, MM and LMW were resolved in 12% gel and stained with coomassie brilliant blue stain. After staining, gel was completely destained and kept in miliQ for overnight. Then, stained protein gel pieces were excised and stain was removed from gel pieces. Gel pieces were reduced with 10 mM DTT followed by alkylation with 50 mM IAA (Iodoacetamide). Then gel pieces were washed and incubated with trypsin (20 μg) and Glu-C (10 μg), separately at 37 ͦ C for overnight. Next, digested peptides were separated from the gel pieces and dried under vacuum. Samples were zip tipped and analysed in MS.

### Proteomics analysis of BV2 cells exposed to different species of α-synuclein Sample preparation

Modified and unmodified α-synuclein treated BV2 cells were harvested and lysed in lysis buffer (8 M urea) and centrifuged to clear cell debris. The supernatant was collected and total protein concentration was determined using Micro BCA™ protein assay kit (Thermo Scientific). Protein samples were then reduced with 10 mM DTT for 1 h at 60 °C and subsequently alkylated using a 20 mM IAA in dark at room temperature for 30 min. The alkylated proteins were then digested by adding sequencing grade trypsin (1:20 w/w in 50 mM TEAB) and incubating at 37 °C for overnight. Digestion was stopped by the addition of formic acid to a final concentration of 0.1 %. The digested solution was passed through a C18 tip to remove salts and buffer ions. The peptides were eluted by 80% ACN with 0.1% formic acid. The eluted samples were vacuum dried and stored at -20 °C for mass spectrometry analysis.

### Quantitative proteomic analysis using Data-Independent acquisition (SWATH-MS)

The digested peptides (50µg) were dissolved in solvent A (98% H2O, 2% acetonitrile with 0.1% formic acid) and each sample was spiked with iRT reagent as recommended by the manufacturer. To collect data in SWATH acquisition mode, the instrument was configured as described by Gillet et al.19. Briefly, the mass spectrometer was operated in a looped product ion mode. Using variable isolation with 1 Da window overlaps, a set of 50 overlapping windows were constructed covering the mass range 400–1200 Da.

Each sample (4 µg peptides in solvent A) was injected in triplicates and data were acquired on Triple TOF 5600+, (Sciex, Concord Canada) which was coupled with a ChromXP reverse-phase 3 μm C18-CL trap column (350 μm × 0.5 mm, 120 Å, Eksigent) and nanoViper C18 separation column (75 μm × 250 mm, 3 μm, 100 Å; Acclaim Pep Map, ThermoScientific) in Eksigent NanoLC-2DPlus system (Eksigent, Dublin, CA, USA). The samples were loaded at a flow rate of 2 µl/min for 10 min using solvent A and peptides were eluted from the analytical column at a flow rate of 300 nl/min in a linear gradient of 5% solvent B to 28% solvent B in 110min with total run time of 120 min. Solvent A was composed of 0.1% (v/v) formic acid in water and solvent B was comprised of 95% (v/v) acetonitrile with 0.1% (v/v) formic acid.

For SWATH-MS data processing and quantification, the mass spectrometry raw files were processed in Spectronaut Pulsar 14 (Biognosys) in the Direct-DIA method using default parameters, data were searched against UniProtKB protein database (contains 20069 mouse protein entries) includes iRT peptides (Biognosys). The FDR was set to 1% at the peptide precursor level. The extracted-ion peak intensities were further used for the quantitative analysis of proteins. T-test was used to compare natural log-transformed protein abundance values of control BV2 cells and treatment conditions including BV2 cells with LMW species of *α*-synuclein (LMW), MGO modified *α*-synuclein LMW (MM), *α*-synuclein oligomers (SO), MGO modified *α*-synuclein oligomers (MO). The proteins were considered to have changed significantly only. Proteins were considered only if they have at least two unique peptides and were significant (abundance changed > 1.3 fold with <0.05 p-value) in all independent biological and technical triplicate experiments. The average value of all biological replicates was used to indicate the final protein abundance at a given condition. The mass spectrometry proteomics data have been deposited to the PRIDE Archive (http://www.ebi.ac.uk/pride/archive/) via the PRIDE partner repository with the data set identifier PXD030349. Data were analyzed using R software packages and GraphPad Prism version 5.02 (GraphPad Software, La Jolla, CA).

### Pathway and Functional enrichment analysis

Reactome pathway term enrichment analysis was performed by g:Profiler (https://biit.cs.ut.ee/gprofiler/) and GSEA v4.1.0 (http://www.broadin-stitute.org/gsea/index.jsp), Cytoscape v3.8.2 with the Enrichment Map Plugin was used for interpretation and visualization of the GSEA and gProfiler results. Genes were ranked from most up-regulated to most down-regulated by using GSEAPreranked function. This pre-ranked list was used for pathway analysis using gene-set enrichment analysis (GSEA). Gene sets used included REACTOME pathway database. For GSEA FDR < 0.1 applied considered statistically significant. Only gene sets consisting of more than 5 and fewer than 500 genes were taken into account. The enrich-ment map was generated with enriched gene sets that passed the significance threshold of P<0.05 and FDR Q<0.05. The enrichment map was summarized by collapsing node clusters and these clusters of nodes were labelled using the Cytoscape AutoAnnotate plug-in. Clusters were manually arranged for clarity.

### Cell culture

Authenticated BV2 cell line (immortalized murine neonatal microglia) were grown in DMEM HiGlutaXL (AL-007G-500 ml, Himedia) media supplemented with 10% FBS (RM9955, Himedia) and maintained at 37 ^0^C along with 5% CO2 humidified atmosphere. HEK-Blue™-hTLR2 cell line (Co-transfected with human TLR2 and SEAP reported gene) was purchased from InvivoGen. These cells were grown in DMEM media supplemented with 10% FBS, HEK- Blue™-selection reagent (hb-sel, InvivoGen) and normocin (50mg/ml) (ant-nr-1, InvivoGen) in 5% CO2 humidified atmosphere at 37 ^0^C.

### Confocal imaging

For activation and species internalization analysis, BV2 cells were grown on coverslips in 6well plate for overnight in DMEM GlutMAX media. Before 4h of treatment, cells were shifted in incomplete media, subsequently 5μM of species of α-synuclein added to the cells for 6 h. Then, BV2 cells were washed with chilled PBS and fixed overnight with chilled methanol. Post-fixation, cells were washed and blocked with PBSAT for 1h at RT. Cells were probed with rabbit polyclonal α-synuclein antibody (26283, CST) and goat monoclonal IBA antibody (PA518039, Invitrogen) for 1h at RT in humid chamber. After washing, cells were incubated with Alexa-Fluor-488-conjugated anti-rabbit secondary antibody (A11034, Molecular Probes) and Alexa-Fluor-594-conjugated anti-goat secondary anibody (A11058, Invitrogen) respectively, for 1h at RT in humid chamber. Nuclei were stained with DAPI (cat no Sigma) for 5 min and mounted with Prolong gold (P36934, Invitrogen) for microscopic examination. For another set of experiment, BV2 cells were seeded on coverslips in 6 well plate. All species were treated to seeded cells at concentration of 5 μM for 6 h in incomplete media. Post 6 h, cells were washed and fixed with methanol and probed for rabbit α-synuclein primary antibody (26283, CST) and clathrin primary antibody (ab129326 abcam) for 1 h at RT in humid chambers. Cells were washed and incubated with Alexa-Fluor-488-conjugated anti-rabbit secondary antibody (A11034, Molecular Probes) and Alexa-Fluor-568-conjugated anti-goat secondary anibody (ab175474, abcam) for 1h at RT. Cells were washed and stained nucleus with DAPI (D9542, Sigma), subsequently, mounted with Prolong gold (P36934, Invitrogen).

For mitophagy experiment, BV2 cells were seeded over coverslips in 12 well plate and treated with α-synuclein species. Post 12 h treatment, cells were washed and treated with Mitotraker for 10 min. After washing, cells were fixed with chilled methanol for 12h. Next, cells were washed and probed for LC3 primary antibody (ab168831, abcam) for 1 h at room temperature in humid chamber. Treated cells were further probed for Alexa-Fluor-488-conjugated anti-rabbit secondary antibody. For all experiments, respective fluorescence was detected using Leica TCS SP8 laser scanning confocal microscope with 63X oil immersion objective. All samples were imaged with z-stack range from bottom to top of the cell. All samples in each set of experiments were acquired with identical acquisition settings.

For localization of TLR4, we seeded BV2 cells over coverslips in 12 well plate and treated with α-synuclein species. Post 12 h treatment, cells were washed and cells were fixed with chilled methanol for 12h. Next, cells were washed and probed for TLR4 primary antibody (MA5-16216, Invitrogen) and RAB7 primary antibody (D95F2, CST) for 1 h at room temperature in humid chamber. Treated cells were further probed for Alexa-Fluor-594- conjugated anti-mouse secondary antibody (A11005, Invitrogen) and Alexa-Fluor-488- conjugated anti-rabbit secondary antibody, respectively. For all experiments, respective fluorescence was detected using Leica TCS SP8 laser scanning confocal microscope with 63X oil immersion objective. All samples were imaged with z-stack range from bottom to top of the cell. All samples in each set of experiments were acquired with identical acquisition settings.

### Confocal image analysis

For all the set of confocal experiments, we have utilized the Leica TCS SP8 analysis software. All images were processed with the confined image settings. Fluorescence intensities were measured after background subtraction, by selecting multiple ROIs in median planes, which include both internalized and membrane bound fluorescence signals. For few experimental, we have constructed 3D image using the z-stack range and Gaussian filter with sigma value 1 was used to process 3D images in Leica TCS SP8 3D software. For colocalization analysis, we have used FIJI (ImageJ) software (NIH, Bethesda, MD) with CoLoc2 plugin. Pearson’s correlation coefficient value above threshold was reported in order to represent colocalization.

### Low force mode atomic force microscopy

Microglia cells (BV2) were grown over coverslips and maintained in DMEM GlutMax, supplemented with 10% FBS. Before treatment, cells were starved for 4 h and treated with 5 μM of α-synuclein species for 12h. Methanol fixed cells were washed with water and imaging of cells was carried out by JPK Nano Wizard III atomic force microscope (JPK instrument, Berlin, Germany). All images were taken in contact mode using gold-coated Hydra cantilever (resonance frequency of 17 kHz (±4 kHz), force constant of 0.1 N/m). All images were taken at scan speed of 0.8-1 line/s with resolution of 256 x 256 and 512 x 512 pixels. Images are processed using JPK process software.

### Western blotting

BV2 cells were cultured in DMEM GlutaMAX media supplemented with 10% fetal bovine serum (FBS), 1% antibiotic in a humidified 5% CO2 atmosphere at 37°C. Cells were treated with α-synuclein species (5 μM) in incomplete media and harvested at different time intervals (6 h and 12 h). Cells were lysed in RIPA buffer (Sigma) and spun at 13k g for 30 min at 4 C. Supernatant were collected and protein estimation were done via BCA method. Protein (30 μg) sample were resolved using 12% SDS-PAGE and transferred to PVDF membrane (MDI, India). Then, membrane were blocked with 5% skimmed milk for 2 h and incubated with RAB7 (D95F2, CST), LC3 (ab168831, abcam), Drp1 (5391P, CST), Mfn2 (11925T, CST), Tom20 (42406T, CST), Bax (2772S, CST), Cytochrome C (PAA594Hu01, SantaCruz), NLRP3 (15101S, CST), Caspase1 (24232S, CST), Myd88 (3699S, CST), TIRAP (13077S, CST), GAPDH primary antibody (1:1000 dilutions, CST) for overnight at 4 ͦ C. The membranes were washed 3 times with PBST before incubated with anti-rabbit secondary antibody (31460, Invitrogen) for 1 h at room temperature. Membrane were probed with chemiluminescent HRP substrate (Luminata) and imaged at ImageQuant LAS 4000 (GE Healthcare).

### Cytokines release assay (ELISA)

To estimate the IL-1β secretion in supernatant, we treated BV2 cells with α-synuclein species (5 μM each) for 24h in incomplete media. The supernatant was collected and centrifuged at 12k rpm for 30 min to remove the cell debris. Then, supernatants were stored in -80 ͦ until analysed by ELISA (Cloud-Clone corp.). Cytokines (IL-1β) were measured as per the manufacture’s protocol. Measurements were done in triplicate and average values were reported.

### SEAP detection HEK TLR2 assay

HEK-Blue cells expressing TLR2 and SEAP reporter gene were seeded into 96 well plate in SEAP detection media (HEK-Blue™ Detection, hb-det3) and immediately treated with 5 μM concentration of α-synuclein species. For this experiment, we used PAM3CSK4 (tlrl-pms, InvivoGen) as a positive control at concentration of 1 μg/ml. After 12 h of treatment, the levels of SEAP in 96 well plate were accessed by monitoring its absorbance at 620 nm. SEAP level is reported as average of three biologicals.

### Statistical analysis

All data were analysed using GraphPad Prism 7.0 software and expressed as mean with standard error of mean (SEM). For comparison between the all groups, ANOVA followed by Tukey’s multiple comparison test were used.

### Availability of data and materials

The proteomics data is available online through the ProteomeXchange Consortium via the PRIDE (https://www.ebi.ac.uk/pride/) partner repository with the dataset identifier PXD030349 for SWATH-MS dataset. All data generated or analyzed during this study are included in this article and the supplementary information.

## Acknowledgements

We thank RCB and ATPC for the technical assistance regarding biophysical and microscopy facilities. We would like to thank to Dr Shubra for a kind support in BLI experiment.

## Authors affiliations

Manisha Kumari, Bhoj Kumar, Krishna Singh Bisht and Tushar Kanti Maiti Functional Proteomics Laboratory, Regional Centre for Biotechnology (RCB), NCR Biotech Science Cluster, 3^rd^ Milestone Gurgaon-Faridabad Expressway, Faridabad, 121001, India.

Bhoj Kumar (Present address)

Complex Carbohydrate Research Center, The University of Georgia, 315 Riverbend Road, Athens, GA 30602, USA

*Corresponding Author: Tushar Kanti Maiti

## Authors’ contributions

MK and TKM designed the study. MK and KSB performed biophysical and cellular experiments. BK and MK designed and executed the mass spectrometry experiments. B.K made substantial contributions to acquisition, interpretation and writing of MS data. MK and TKM analysed the data and wrote the manuscript. All authors read and approved the final manuscript.

## References

1. Ascherio A, Schwarzschild MA. The epidemiology of Parkinson’s disease: risk factors and prevention. The Lancet Neurology. 2016 Nov 1;15(12):1257–72.

2. Werner P, Seppi K, Tanner CM, Halliday GM, Brundin P, Volkmann J, Schrag AE, Lang AE. Parkinson disease. Nat. Rev. Dis. Primers. 2017;3(17013).

3. Burré J, Sharma M, Tsetsenis T, Buchman V, Etherton MR, Südhof TC. α-Synuclein promotes SNARE-complex assembly in vivo and in vitro. Science. 2010 Sep 24;329(5999):1663–7.

4. Janezic S, Threlfell S, Dodson PD, Dowie MJ, Taylor TN, Potgieter D, Parkkinen L, Senior SL, Anwar S, Ryan B, Deltheil T. Deficits in dopaminergic transmission precede neuron loss and dysfunction in a new Parkinson model. Proceedings of the National Academy of Sciences. 2013 Oct 15;110(42):E4016–25.

5. Zasso J, Ahmed M, Cutarelli A, Conti L. Inducible alpha-synuclein expression affects human neural stem cells’ behavior. Stem cells and development. 2018 Jul 15;27(14):985–94.

6. Jin H, Kanthasamy A, Ghosh A, Yang Y, Anantharam V, Kanthasamy AG. α-Synuclein negatively regulates protein kinase Cδ expression to suppress apoptosis in dopaminergic neurons by reducing p300 histone acetyltransferase activity. Journal of Neuroscience. 2011 Feb 9;31(6):2035–51.

7. Perez RG, Waymire JC, Lin E, Liu JJ, Guo F, Zigmond MJ. A role for α-synuclein in the regulation of dopamine biosynthesis. Journal of Neuroscience. 2002 Apr 15;22(8):3090–9.

8. Eliezer D, Kutluay E, Bussell Jr R, Browne G. Conformational properties of α-synuclein in its free and lipid-associated states. Journal of molecular biology. 2001 Apr 6;307(4):1061–73.

9. Giasson BI, Murray IV, Trojanowski JQ, Lee VM. A hydrophobic stretch of 12 amino acid residues in the middle of α-synuclein is essential for filament assembly. Journal of Biological Chemistry. 2001 Jan 26;276(4):2380–6.

10. Stephens AD, Zacharopoulou M, Schierle GS. The cellular environment affects monomeric α-synuclein structure. Trends in biochemical sciences. 2019 May 1;44(5):453–66.

11. Candelise N, Schmitz M, Thüne K, Cramm M, Rabano A, Zafar S, Stoops E, Vanderstichele H, Villar-Pique A, Llorens F, Zerr I. Effect of the micro-environment on α- synuclein conversion and implication in seeded conversion assays. Translational neurodegeneration. 2020 Dec;9(1):1–6.

12. Fujiwara H, Hasegawa M, Dohmae N, Kawashima A, Masliah E, Goldberg MS, Shen J, Takio K, Iwatsubo T. α-Synuclein is phosphorylated in synucleinopathy lesions. Nature cell biology. 2002 Feb;4(2):160–4.

13. Marotta NP, Lin YH, Lewis YE, Ambroso MR, Zaro BW, Roth MT, Arnold DB, Langen R, Pratt MR. O-GlcNAc modification blocks the aggregation and toxicity of the protein α- synuclein associated with Parkinson’s disease. Nature chemistry. 2015 Nov;7(11):913–20.

14. Burai R, Ait-Bouziad N, Chiki A, Lashuel HA. Elucidating the role of site-specific nitration of α-synuclein in the pathogenesis of Parkinson’s disease via protein semisynthesis and mutagenesis. Journal of the American Chemical Society. 2015 Apr 22;137(15):5041–52.

15. Fauvet B, Fares MB, Samuel F, Dikiy I, Tandon A, Eliezer D, Lashuel HA. Characterization of semisynthetic and naturally Nα-acetylated α-synuclein in vitro and in intact cells: implications for aggregation and cellular properties of α-synuclein. Journal of Biological Chemistry. 2012 Aug 17;287(34):28243–62.

16. Kalia LV, Kalia SK, Chau H, Lozano AM, Hyman BT, McLean PJ. Ubiquitinylation of α- synuclein by carboxyl terminus Hsp70-interacting protein (CHIP) is regulated by Bcl-2- associated athanogene 5 (BAG5). PloS one. 2011 Feb 16;6(2):e14695.

17. Rott R, Szargel R, Shani V, Hamza H, Savyon M, Abd Elghani F, Bandopadhyay R, Engelender S. SUMOylation and ubiquitination reciprocally regulate α-synuclein degradation and pathological aggregation. Proceedings of the National Academy of Sciences. 2017 Dec 12;114(50):13176–81.

18. Shaikh S, Nicholson LF. Advanced glycation end products induce in vitro cross-linking of α-synuclein and accelerate the process of intracellular inclusion body formation. Journal of neuroscience research. 2008 Jul;86(9):2071–82.

19. Hassan A, Kandel RS, Mishra R, Gautam J, Alaref A, Jahan N. Diabetes mellitus and Parkinson’s disease: shared pathophysiological links and possible therapeutic implications. Cureus. 2020 Aug;12(8).

20. Yao D, Brownlee M. Hyperglycemia-induced reactive oxygen species increase expression of the receptor for advanced glycation end products (RAGE) and RAGE ligands. Diabetes. 2010 Jan 1;59(1):249–55.

21. Allaman I, Bélanger M, Magistretti PJ. Methylglyoxal, the dark side of glycolysis. Frontiers in neuroscience. 2015 Feb 9;9:23.

22. Candido R, Forbes JM, Thomas MC, Thallas V, Dean RG, Burns WC, Tikellis C, Ritchie RH, Twigg SM, Cooper ME, Burrell LM. A breaker of advanced glycation end products attenuates diabetes-induced myocardial structural changes. Circulation research. 2003 Apr 18;92(7):785–92.

23. Vicente Miranda H, Szegő ÉM, Oliveira LM, Breda C, Darendelioglu E, de Oliveira RM, Ferreira DG, Gomes MA, Rott R, Oliveira M, Munari F. Glycation potentiates α-synuclein-associated neurodegeneration in synucleinopathies. Brain. 2017 May 1;140(5):1399–419.

24. Martinez-Orozco H, Marino L, Uceda AB, Ortega-Castro J, Vilanova B, Frau J, Adrover M. Nitration and Glycation Diminish the α-Synuclein Role in the Formation and Scavenging of Cu2+-Catalyzed Reactive Oxygen Species. ACS chemical neuroscience. 2019 Apr 11;10(6):2919–30.

25. Sharma N, Rao SP, Kalivendi SV. The deglycase activity of DJ-1 mitigates α-synuclein glycation and aggregation in dopaminergic cells: Role of oxidative stress mediated downregulation of DJ-1 in Parkinson’s disease. Free Radical Biology and Medicine. 2019 May 1;135:28–37.

26. Vicente Miranda H, Chegão A, Oliveira MS, Fernandes Gomes B, Enguita FJ, Outeiro TF. Hsp27 reduces glycation-induced toxicity and aggregation of alpha-synuclein. The FASEB Journal. 2020 May;34(5):6718–28.

27. Leake I. Specific pattern of microglial activation in PD. Nature Reviews Neurology. 2021 Jan;17(1):2–2.

28. Imamura K, Hishikawa N, Sawada M, Nagatsu T, Yoshida M, Hashizume Y. Distribution of major histocompatibility complex class II-positive microglia and cytokine profile of Parkinson’s disease brains. Acta neuropathologica. 2003 Dec 1;106(6):518–26.

29. Kim YS, Joh TH. Microglia, major player in the brain inflammation: their roles in the pathogenesis of Parkinson’s disease. Experimental & molecular medicine. 2006 Aug;38(4):333–47.

30. Choi I, Zhang Y, Seegobin SP, Pruvost M, Wang Q, Purtell K, Zhang B, Yue Z. Microglia clear neuron-released α-synuclein via selective autophagy and prevent neurodegeneration. Nature communications. 2020 Mar 13;11(1):1–4.

31. George S, Rey NL, Tyson T, Esquibel C, Meyerdirk L, Schulz E, Pierce S, Burmeister AR, Madaj Z, Steiner JA, Escobar Galvis ML. Microglia affect α-synuclein cell-to-cell transfer in a mouse model of Parkinson’s disease. Molecular neurodegeneration. 2019 Dec;14(1):1–22.

32. Xia Y, Zhang G, Kou L, Yin S, Han C, Hu J, Wan F, Sun Y, Wu J, Li Y, Huang J. Reactive microglia enhance the transmission of exosomal alpha-synuclein via toll-like receptor 2. Brain. 2021 Apr 1.

33. Hughes CD, Choi ML, Ryten M, Hopkins L, Drews A, Botía JA, Iljina M, Rodrigues M, Gagliano SA, Gandhi S, Bryant C. Picomolar concentrations of oligomeric alpha-synuclein sensitizes TLR4 to play an initiating role in Parkinson’s disease pathogenesis. Acta neuropathologica. 2019 Jan;137(1):103–20.

34. Hou L, Bao X, Zang C, Yang H, Sun F, Che Y, Wu X, Li S, Zhang D, Wang Q. Integrin CD11b mediates α-synuclein-induced activation of NADPH oxidase through a Rho-dependent pathway. Redox biology. 2018 Apr 1;14:600–8.

35. Cao S, Standaert DG, Harms AS. The gamma chain subunit of Fc receptors is required for alpha-synuclein-induced pro-inflammatory signaling in microglia. Journal of neuroinflammation. 2012 Dec;9(1):1–1.

36. Kim C, Ho DH, Suk JE, You S, Michael S, Kang J, Lee SJ, Masliah E, Hwang D, Lee HJ, Lee SJ. Neuron-released oligomeric α-synuclein is an endogenous agonist of TLR2 for paracrine activation of microglia. Nature communications. 2013 Mar 5;4(1):1–2.

37. Fan Z, Pan YT, Zhang ZY, Yang H, Yu SY, Zheng Y, Ma JH, Wang XM. Systemic activation of NLRP3 inflammasome and plasma α-synuclein levels are correlated with motor severity and progression in Parkinson’s disease. Journal of neuroinflammation. 2020 Dec;17(1):1–0.

38. von Herrmann KM, Salas LA, Martinez EM, Young AL, Howard JM, Feldman MS, Christensen BC, Wilkins OM, Lee SL, Hickey WF, Havrda MC. NLRP3 expression in mesencephalic neurons and characterization of a rare NLRP3 polymorphism associated with decreased risk of Parkinson’s disease. NPJ Parkinson’s disease. 2018 Aug 15;4(1):1–9.

39. Anderson FL, von Herrmann KM, Andrew AS, Kuras YI, Young AL, Scherzer CR, Hickey WF, Lee SL, Havrda MC. Plasma-borne indicators of inflammasome activity in Parkinson’s disease patients. NPJ Parkinson’s disease. 2021 Jan 4;7(1):1–2.

40. Paleologou KE, Schmid AW, Rospigliosi CC, Kim HY, Lamberto GR, Fredenburg RA, Lansbury PT, Fernandez CO, Eliezer D, Zweckstetter M, Lashuel HA. Phosphorylation at Ser- 129 but not the phosphomimics S129E/D inhibits the fibrillation of α-synuclein. Journal of Biological Chemistry. 2008 Jun 13;283(24):16895–905.

41. Volles MJ, Lansbury Jr PT. Relationships between the sequence of α-synuclein and its membrane affinity, fibrillization propensity, and yeast toxicity. Journal of molecular biology. 2007 Mar 9;366(5):1510–22.

42. Ghosh D, Maji SK. Preparation of aggregate-free α-synuclein for in vitro aggregation study. Protoc. Exchange. 2015 Apr 18:270–81.

43. Kumari P, Ghosh D, Vanas A, Fleischmann Y, Wiegand T, Jeschke G, Riek R, Eichmann C. Structural insights into α-synuclein monomer–fibril interactions. Proceedings of the National Academy of Sciences. 2021 Mar 9;118(10).

44. Giasson BI, Murray IV, Trojanowski JQ, Lee VM. A hydrophobic stretch of 12 amino acid residues in the middle of α-synuclein is essential for filament assembly. Journal of Biological Chemistry. 2001 Jan 26;276(4):2380–6.

45. Stöckl MT, Zijlstra N, Subramaniam V. α-Synuclein oligomers: an amyloid pore?. Molecular neurobiology. 2013 Apr;47(2):613–21.

46. Wendeln AC, Degenhardt K, Kaurani L, Gertig M, Ulas T, Jain G, Wagner J, Häsler LM, Wild K, Skodras A, Blank T. Innate immune memory in the brain shapes neurological disease hallmarks. Nature. 2018 Apr;556(7701):332-8.

47. Gheorghe RO, Deftu A, Filippi A, Grosu A, Bica-Popi M, Chiritoiu M, Chiritoiu G, Munteanu C, Silvestro L, Ristoiu V. Silencing the cytoskeleton protein Iba1 (ionized calcium binding adapter protein 1) interferes with BV2 microglia functioning. Cellular and molecular neurobiology. 2020 Jan 16:1–7.

48. Ito D, Tanaka K, Suzuki S, Dembo T, Fukuuchi Y. Enhanced expression of Iba1, ionized calcium-binding adapter molecule 1, after transient focal cerebral ischemia in rat brain. Stroke. 2001 May;32(5):1208–15.

49. Caldeira C, Oliveira AF, Cunha C, Vaz AR, Falcão AS, Fernandes A, Brites D. Microglia change from a reactive to an age-like phenotype with the time in culture. Frontiers in cellular neuroscience. 2014 Jun 2;8:152.

50. Saraswathy S, Wu G, Rao NA. Retinal microglial activation and chemotaxis by docosahexaenoic acid hydroperoxide. Investigative ophthalmology & visual science. 2006 Aug 1;47(8):3656–63.

51. Lee HJ, Suk JE, Bae EJ, Lee JH, Paik SR, Lee SJ. Assembly-dependent endocytosis and clearance of extracellular a-synuclein. The international journal of biochemistry & cell biology. 2008 Jan 1;40(9):1835–49.

52. Fortin DL, Troyer MD, Nakamura K, Kubo SI, Anthony MD, Edwards RH. Lipid rafts mediate the synaptic localization of α-synuclein. Journal of Neuroscience. 2004 Jul 28;24(30):6715–23.

53. Ben Gedalya T, Loeb V, Israeli E, Altschuler Y, Selkoe DJ, Sharon R. α-Synuclein and Polyunsaturated Fatty Acids Promote Clathrin-Mediated Endocytosis and Synaptic Vesicle Recycling. Traffic. 2009 Feb;10(2):218–34.

54. Kaksonen M, Roux A. Mechanisms of clathrin-mediated endocytosis. Nature reviews Molecular cell biology. 2018 May;19(5):313–26.

55. Schuh AL, Audhya A. The ESCRT machinery: from the plasma membrane to endosomes and back again. Critical reviews in biochemistry and molecular biology. 2014 May 1;49(3):242–61.

56. Park JS, Davis RL, Sue CM. Mitochondrial dysfunction in Parkinson’s disease: new mechanistic insights and therapeutic perspectives. Current neurology and neuroscience reports. 2018 May;18(5):1–1.

57. Smits WK, Dubois JY, Bron S, van Dijl JM, Kuipers OP. Tricksy business: transcriptome analysis reveals the involvement of thioredoxin A in redox homeostasis, oxidative stress, sulfur metabolism, and cellular differentiation in Bacillus subtilis. Journal of bacteriology. 2005 Jun 15;187(12):3921–30.

58. Guo JD, Zhao X, Li Y, Li GR, Liu XL. Damage to dopaminergic neurons by oxidative stress in Parkinson’s disease. International journal of molecular medicine. 2018 Apr 1;41(4):1817–25.

59. Hoshino A, Wang WJ, Wada S, McDermott-Roe C, Evans CS, Gosis B, Morley MP, Rathi KS, Li J, Li K, Yang S. The ADP/ATP translocase drives mitophagy independent of nucleotide exchange. Nature. 2019 Nov;575(7782):375–9.

60. Runwal G, Stamatakou E, Siddiqi FH, Puri C, Zhu Y, Rubinsztein DC. LC3-positive structures are prominent in autophagy-deficient cells. Scientific reports. 2019 Jul 12;9(1):1–4.

61. Roodveldt C, Labrador-Garrido A, Gonzalez-Rey E, Lachaud CC, Guilliams T, Fernandez- Montesinos R, Benitez-Rondan A, Robledo G, Hmadcha A, Delgado M, Dobson CM. Preconditioning of microglia by α-synuclein strongly affects the response induced by toll-like receptor (TLR) stimulation. PloS one. 2013 Nov 13;8(11):e79160.

62. Daniele SG, Béraud D, Davenport C, Cheng K, Yin H, Maguire-Zeiss KA. Activation of MyD88-dependent TLR1/2 signaling by misfolded α-synuclein, a protein linked to neurodegenerative disorders. Science signaling. 2015 May 12;8(376):ra45.

63. Pascual-Lucas M, Fernandez-Lizarbe S, Montesinos J, Guerri C. LPS or ethanol triggers clathrin-and rafts/caveolae-dependent endocytosis of TLR 4 in cortical astrocytes. Journal of neurochemistry. 2014 May;129(3):448–62.

64. Husebye H, Halaas Ø, Stenmark H, Tunheim G, Sandanger Ø, Bogen B, Brech A, Latz E, Espevik T. Endocytic pathways regulate Toll-like receptor 4 signaling and link innate and adaptive immunity. The EMBO journal. 2006 Feb 22;25(4):683–92.

65. Mishra SR, Mahapatra KK, Behera BP, Patra S, Bhol CS, Panigrahi DP, Praharaj PP, Singh A, Patil S, Dhiman R, Bhutia SK. Mitochondrial dysfunction as a driver of NLRP3 inflammasome activation and its modulation through mitophagy for potential therapeutics. The International Journal of Biochemistry & Cell Biology. 2021 Jul 1;136:106013.

66. Abais JM, Xia M, Zhang Y, Boini KM, Li PL. Redox regulation of NLRP3 inflammasomes: ROS as trigger or effector?. Antioxidants & redox signaling. 2015 May 1;22(13):1111–29.

67. Murakami T, Ockinger J, Yu J, Byles V, McColl A, Hofer AM, Horng T. Critical role for calcium mobilization in activation of the NLRP3 inflammasome. Proceedings of the National Academy of Sciences. 2012 Jul 10;109(28):11282–7.

68. Bousset L, Pieri L, Ruiz-Arlandis G, Gath J, Jensen PH, Habenstein B, Madiona K, Olieric V, Böckmann A, Meier BH, Melki R. Structural and functional characterization of two alpha- synuclein strains. Nature communications. 2013 Oct 10;4:2575.

69. Ingelsson M. Alpha-Synuclein Oligomers—Neurotoxic Molecules in Parkinson’s Disease and Other Lewy Body Disorders. Frontiers in neuroscience. 2016;10.

70. Chen SW, Drakulic S, Deas E, Ouberai M, Aprile FA, Arranz R, Ness S, Roodveldt C, Guilliams T, De-Genst EJ, Klenerman D. Structural characterization of toxic oligomers that are kinetically trapped during α-synuclein fibril formation. Proceedings of the National Academy of Sciences. 2015 Apr 21;112(16):E1994–2003.

71. Chakraborty, R., Dey, S., Sil, P. et al. Conformational distortion in a fibril-forming oligomer arrests alpha-Synuclein fibrillation and minimizes its toxic effects. Commun Biol 4, 518 (2021).

72. Elkin SR, Lakoduk AM, Schmid SL. Endocytic pathways and endosomal trafficking: a primer. Wiener Medizinische Wochenschrift. 2016 May;166(7):196–204.

73. Krzystek TJ, Banerjee R, Thurston L, Huang J, Swinter K, Rahman SN, Falzone TL, Gunawardena S. Differential mitochondrial roles for α-synuclein in DRP1-dependent fission and PINK1/Parkin-mediated oxidation. Cell death & disease. 2021 Aug 17;12(9):1–6.

74. Di Maio R, Barrett PJ, Hoffman EK, Barrett CW, Zharikov A, Borah A, Hu X, McCoy J, Chu CT, Burton EA, Hastings TG. α-Synuclein binds to TOM20 and inhibits mitochondrial protein import in Parkinson’s disease. Science translational medicine. 2016 Jun 8;8(342):342ra78-.

75. Williams JA, Ding WX. Mechanisms, pathophysiological roles and methods for analyzing mitophagy–recent insights. Biological chemistry. 2018 Feb 1;399(2):147–78.

76. Grassi D, Howard S, Zhou M, Diaz-Perez N, Urban NT, Guerrero-Given D, Kamasawa N, Volpicelli-Daley LA, LoGrasso P, Lasmézas CI. Identification of a highly neurotoxic α- synuclein species inducing mitochondrial damage and mitophagy in Parkinson’s disease. Proceedings of the National Academy of Sciences. 2018 Mar 13;115(11):E2634–43.

77. Pike AF, Varanita T, Herrebout MA, Plug BC, Kole J, Musters RJ, Teunissen CE, Hoozemans JJ, Bubacco L, Veerhuis R. α-Synuclein evokes NLRP3 inflammasome-mediated IL-1β secretion from primary human microglia. Glia. 2021 Jun;69(6):1413–28.

78. Panicker N, Sarkar S, Harischandra DS, Neal M, Kam TI, Jin H, Saminathan H, Langley M, Charli A, Samidurai M, Rokad D. Fyn kinase regulates misfolded α-synuclein uptake and NLRP3 inflammasome activation in microglia. The Journal of experimental medicine. 2019 Jun 3;216(6):1411–30.

79. Zhong Z, Umemura A, Sanchez-Lopez E, Liang S, Shalapour S, Wong J, He F, Boassa D, Perkins G, Ali SR, McGeough MD. NF-κB restricts inflammasome activation via elimination of damaged mitochondria. Cell. 2016 Feb 25;164(5):896–910.

