## Supplementary Figures and Tables for "Glycation renders ɑ-synuclein oligomeric strain and modulates microglia activation"

### Supplementary information

**Supplementary Figure 1:** Mass spectrometry analysis of MGO modified sites in α-synuclein (A) Table representing identified MGO modified lysine residues along with their mass and identification score. (B) Fragmentation of ions in unmodified and modified α-synuclein. (C) MGO modified lysine residues represented with M in the sequence of α-synuclein.

**Supplementary Figure 2: α-Synuclein species modulates morphology of BV2 cells.** (A) Atomic force microscopy representing change in morphology of BV2 cells under treatment α-synuclein species. 2D sobel image segregate artefacts from actual AFM image. Scale bar, 20 μm. Subsequently, inset of 2D AFM images showing processes of microglia. Scale bar, 5 μm (B) Confocal images of F-actin stained cells showing modulation of processes of BV2 after treatment α-synculein species.

**Supplementary Figure 3:** Quantification of α-synuclein species internalized in BV2 cells using Mass spectrometry.

**Supplementary Figure 4:** Pathway Enrichment Map visualization of the GSEA analysis.

**Supplementary Figure 5:** Relative protein expression of HSPA8 in different treatment conditions, identified in MS data.

**Supplementary Table 1:** Kinetic parameters of binding of MGO modified species of α-synuclein with lipid membrane.

**Supplementary Table 2:** Kinetic parameters of binding of MGO modified species of α-synuclein with Low molecular weight (LMW) synuclein.

**Supplementary Table 3:** Kinetic parameters of binding of MGO modified species of α-synuclein with TLR2.

**Supplementary Data 1:** List of significantly changed proteins and relative abundance in different pairs of treatment condition.

**Supplementary Data 2:** Gene Set Enrichment Analysis (GSEA) of modulated protein in treatment comparison in pair of MM/LMW, MO/LMW, SO/LMW and MO/SO.

**Supplementary Data 3**: gProfiler enriched Reactome pathway analysis of all modulated genes.

**Supplementary Data 4:** gProfiler enriched Reactome pathway analysis of upregulated genes in MO/SO.

**Supplementary Data 5:** gProfiler enriched Reactome pathway analysis of downregulated genes in MO/SO.

**Supplementary Figure S1**

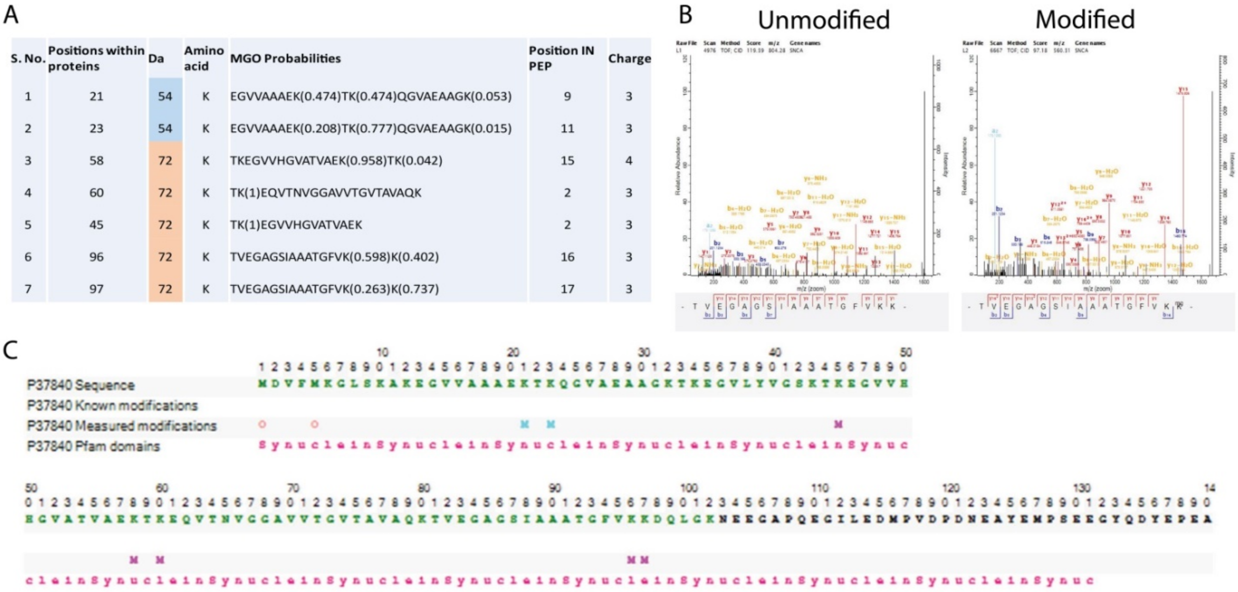

| S. No. | Positions within proteins | Da | Amino acid | MGO Probabilities | Position IN PEP | Charge |
|---|---|---|---|---|---|---|
| 1 | 21 | 54 | K | EGVVAAAEK(0.474)TK(0.474)QGVAEAAGK(0.053) | 9 | 3 |
| 2 | 23 | 54 | K | EGVVAAAEK(0.208)TK(0.777)QGVAEAAGK(0.015) | 11 | 3 |
| 3 | 58 | 72 | K | TKEGVVHGVATVAEK(0.958)TK(0.042) | 15 | 4 |
| 4 | 60 | 72 | K | TK(1)EQVTNVGGAVVTGVTAVAQK | 2 | 3 |
| 5 | 45 | 72 | K | TK(1)EGVVHGVATVAEK | 2 | 3 |
| 6 | 96 | 72 | K | TVEGAGSIAAATGFVK(0.598)K(0.402) | 16 | 3 |
| 7 | 97 | 72 | K | TVEGAGSIAAATGFVK(0.263)K(0.737) | 17 | 3 |



**Supplementary Figure S1: Mass spectrometry analysis of MGO modified sites in α-synuclein**. (A) MGO modification and position on Alpha-synuclein was confirmed by peptide fragmentation using Mass spectrometry, a total 6 position was identified as MGO modified position. (B) MS/MS spectral profile of MGO modified and unmodified peptide. (C) Sequence coverage (green alphabets) of Alpha-synuclein and its MGO modification site donated by M in magenta and pink color for 54 and 72 Da respectively.

### Supplementary Figure S2

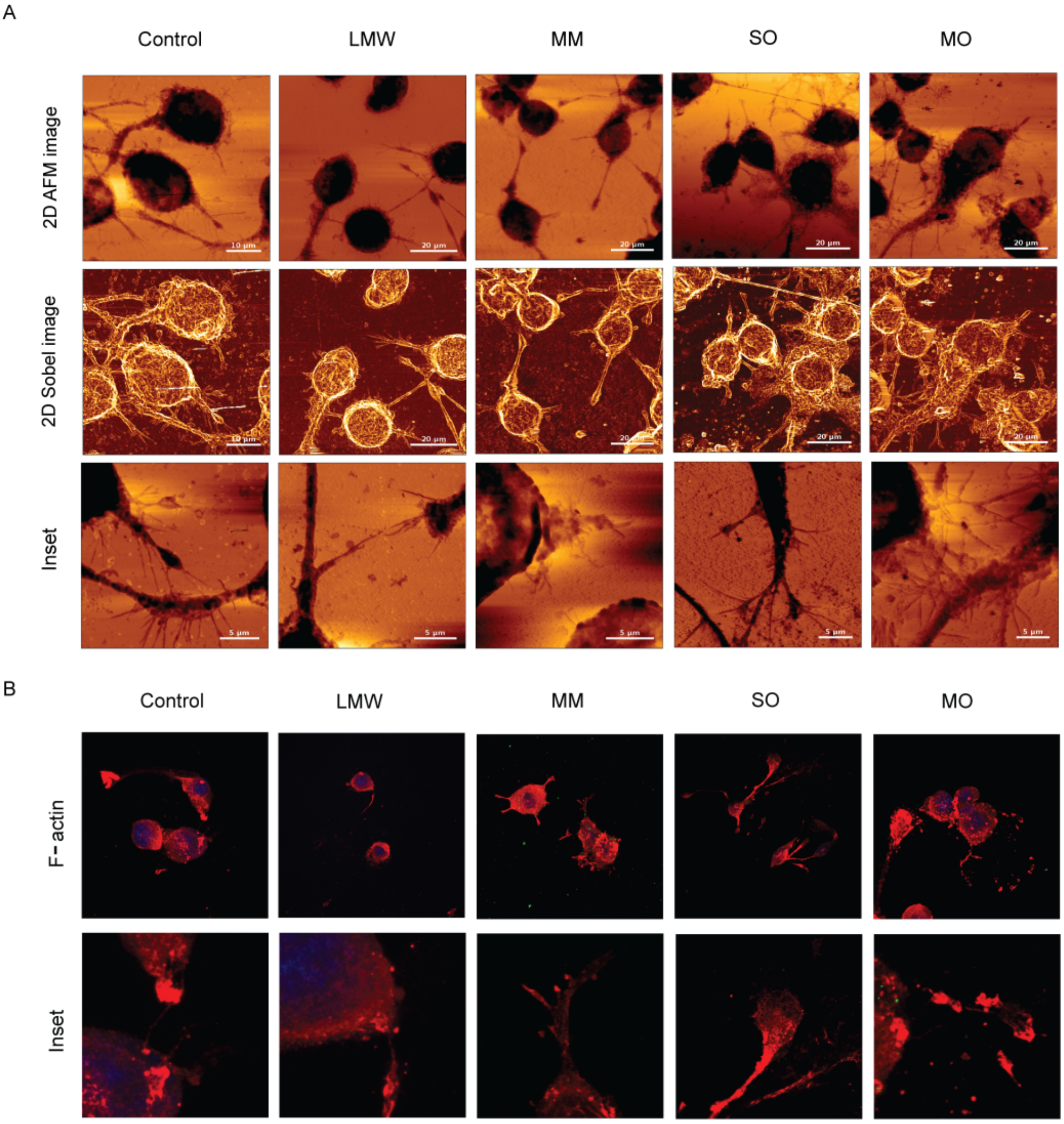


**Supplementary Figure S2: α-Synuclein species modulates morphology of BV2 cells.** (A) Atomic force microscopy representing change in morphology of BV2 cells under treatment α-synuclein species. 2D sobel image segregate artefacts from actual AFM image. Scale bar, 20 µm. Subsequently, inset of 2D AFM images showing processes of microglia. Scale bar, 5 µm (B) Confocal images of F-actin stained cells showing modulation of processes of BV2 after treatment α-synuclein species.

**Supplementary Figure S3**

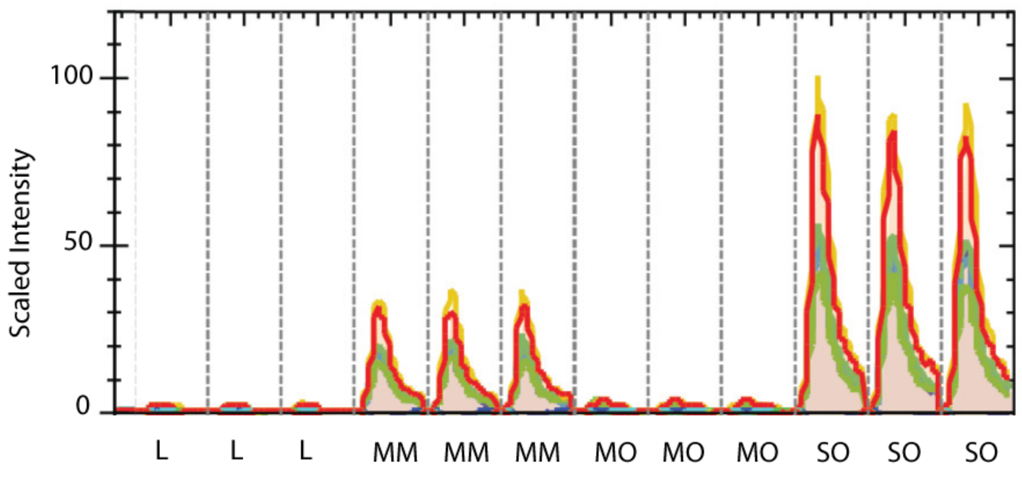


**Supplementary Figure S3:** Quantification of α-synuclein species internalized in BV2 cells using Mass spectrometry.

**Supplementary Figure S4**

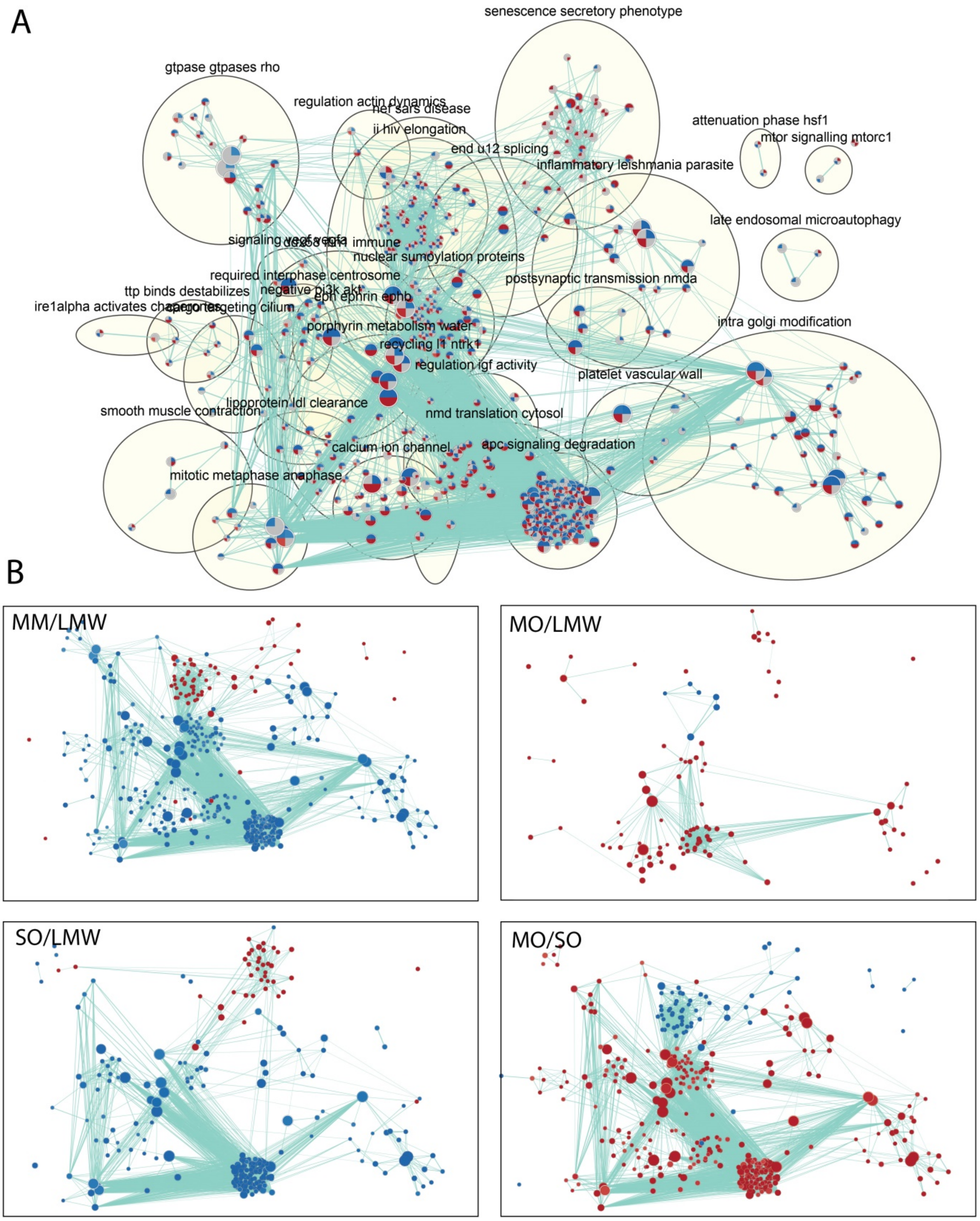


**Supplementary Figure S4: Pathway Enrichment Map visualization of the GSEA analysis.** (A) All modulated genes in MO/SO condition was used for pathway enrichment. (B) Their relative abundance of significance genes were shown in pair of MM/LMW, MO/LMW, SO/LMW and MO/SO. Each node represents enriched Reactome pathway term, node size is proportional to the number of genes in each node. Node color represents the normalized enrichment scores (NES), blue and red color represents down and up-regulated nodes respectively. The thickness of edges represents the richness of shared genes between connected nodes. Edges between nodes overlap of member genes between pathway terms. AutoAnnotate function was applied to group gene-sets those belong similar category and pathways are depicted as circles (A).

**Supplementary Figure S5**

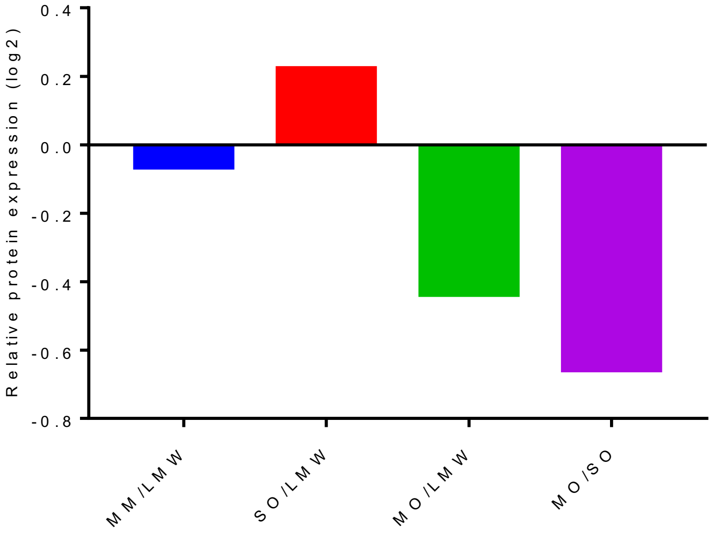


**Supplementary Figure S5:** Relative protein expression of HSPA8 in different treatment conditions, identified in MS data.

**Supplementary Table S1:** Kinetic parameters of binding of MGO modified species of α-synuclein with lipid membrane.

| Samples | kon(1/Ms) | kdis(1/s) | KD (M) | Full R^2 |
|---|---|---|---|---|
| LMW | 2.89E+03±1.5 | 3.38E-03±3.4 | 9.66E-07±5.9 | 0.634067±0.1 |
| MM | 4.51E+02±2.9 | 1.04E+00±0.5 | 2.42E-03±0.31 | 0.9097±0.05 |

**Supplementary Table S2:** Kinetic parameters of binding of MGO modified species of α-synuclein with Low molecular weight (LMW) synuclein.

| Samples | kon(1/Ms) | kdis(1/s) | KD (M) | Full R^2 |
|---|---|---|---|---|
| LMW | 7.26E+06±10 | 1.01E-02±30.2 | 3.74E-08±5.2 | 0.8492±0.01 |
| MM | 1.34E+03±1.8 | 2.41E-03±1.5 | 1.04E-05±1.2 | 0.98715±0.001 |
| SO | 7.72E+04±3.9 | 2.74E-01±2.9 | 2.98E-06±2.2 | 0.99055±0.01 |
| MO | 2.80E+02±2.6 | 1.11E-01±1.5 | 2.44E-04±3.2 | 0.9905±0.004 |

**Supplementary Table S3:** Kinetic parameters of binding of MGO modified species of α-synuclein with TLR2.

| Samples | kon(1/Ms) | kdis(1/s) | KD (M) | Full R^2 |
|---|---|---|---|---|
| LMW | 2.71E+05±2.3 | 1.62E-03±1.0 | 1.50E-07±2.5 | 0.9541±0.03 |
| MM | ND | ND | ND | ND |
| SO | 6.93E+04±6.3 | 3.25E-03±2.7 | 7.16E-08±4.7 | 0.9494±0.03 |
| MO | ND | ND | ND | ND |
| PAM3CSK4 | 8.09E+02±2.0 | 1.68E-03±2.0 | 1.67E-06±2.2 | 0.9206±0.07 |
